# Control of mammalian brain aging by the unfolded protein response (UPR)

**DOI:** 10.1101/2020.04.13.039172

**Authors:** Felipe Cabral-Miranda, Giovani Tamburini, Gabriela Martinez, Danilo Medinas, Yannis Gerakis, Tim Miedema, Claudia Duran-Aniotz, Alvaro O. Ardiles, Cristobal Gonzalez, Carleen Sabusap, Francisca Bermedo-Garcia, Stuart Adamson, Kaitlyn Vitangcol, Hernan Huerta, Xu Zhang, Tomohiro Nakamura, Sergio Pablo Sardi, Stuart A. Lipton, Brian K. Kenedy, Julio Cesar Cárdenas, Adrian G. Palacios, Lars Plate, Juan Pablo Henriquez, Claudio Hetz

**Affiliations:** Biomedical Neuroscience Institute, Faculty of Medicine, University of Chile, Santiago, Chile; Center for Geroscience, Brain Health and Metabolism, Santiago, Chile; Program of Cellular and Molecular Biology, Center for Molecular Studies of the Cell, Institute of Biomedical Sciences, University of Chile, Santiago, Chile; Instituto de Ciências Biomédicas, Universidade do Rio de Janeiro, Rio de Janeiro, Brazil; Centro Interdisciplinario de Neurociencia de Valparaíso, Universidad de Valparaiso, Valparaiso, Chile; Centro de Neurología Traslacional, Escuela de Medicina, Universidad de Valparaíso, Valparaiso, Chile; Department of Chemistry, Vanderbilt University, Nashville, TN, USA; Department of Cell Biology, Center for Advanced Microscopy (CMA BioBio), Universidad de Concepción, Concepción, Chile; Buck Institute for Research on Aging, Novato, CA, USA; Center for Integrative Biology, Universidad Mayor, Santiago, Chile; Department of Molecular Medicine and Neuroscience Translational Center, The Scripps Research Institute, La Jolla, CA 92037, USA; Rare and Neurological Diseases Therapeutic Area, Sanofi, 49 New York Avenue, Framingham, MA, 01701, USA; Department of Neurosciences, University of California, San Diego, School of Medicine, La Jolla, CA 92093, USA; Singapore Institute for Clinical Sciences, Agency for Science, Technology and Research (A*STAR), Singapore

**Author notes:** Correspondence to or. Website: www.hetzlab.cl.

## Abstract

Aging is the major risk factor for the development of dementia and neurodegenerative disorders, and the aging brain manifests severe deficits in buffering capacity by the proteostasis network. Accordingly, we investigated the significance of the unfolded protein response (UPR), a major signaling pathway that copes with endoplasmic reticulum (ER) stress, to normal mammalian brain aging. Genetic disruption of ER stress sensor IRE1α accelerated cognitive and motor dysfunction during aging. Exogenous bolstering of the UPR by overexpressing an active form of the transcription factor XBP1 restored synaptic and cognitive function in addition to reducing cell senescence. Remarkably, proteomic profiling of hippocampal tissue indicated that XBP1s expression corrected age-related alterations in synaptic function. Collectively, our data demonstrate that strategies to manipulate the UPR sustain healthy brain aging.

**One Sentence Summary:** The IRE1/XBP1 pathway dictates when and how brain function declines during aging.

## Main Text

Normal aging is associated with progressive cognitive impairment, representing the most prevalent risk factor for the development of dementia in neurodegenerative disorders. Subtle structural and functional alterations in synapses are the main drivers of age-related cognitive decline (reviewed in *1*), but the molecular mechanisms dictating these perturbations are still elusive. Decades of research have defined the hallmarks of aging, underscoring the biological processes that determine when and how organisms age, thus regulating healthspan and lifespan (*2,3*). Proteostasis (homeostasis of proteins) is maintained by the dynamic integration of pathways that mediate the synthesis, folding, degradation, quality control, trafficking and targeting of proteins, and its disturbance has been posited as a pillar of the aging process (*4*). The complexity of synaptic architecture and its dynamic regulation highlight the need to maintain the integrity of proteostasis at the level of the secretory pathway during the organismal lifespan to sustain normal brain function (*5*).

In order to evaluate and quantify cognitive and motor decline associated with normal aging in rodents, we used a battery of tasks to assess the behavior of young (3 month-old), middle-aged (12 month-old), and aged (18 month-old) mice (Fig. 1A). We detected spontaneous decline in the performance of animals starting at middle age and progressing thereafter using several cognitive and motor evaluations (Fig. 1A and fig. S1), as previously reported *(6)*.

**Fig. 1.**
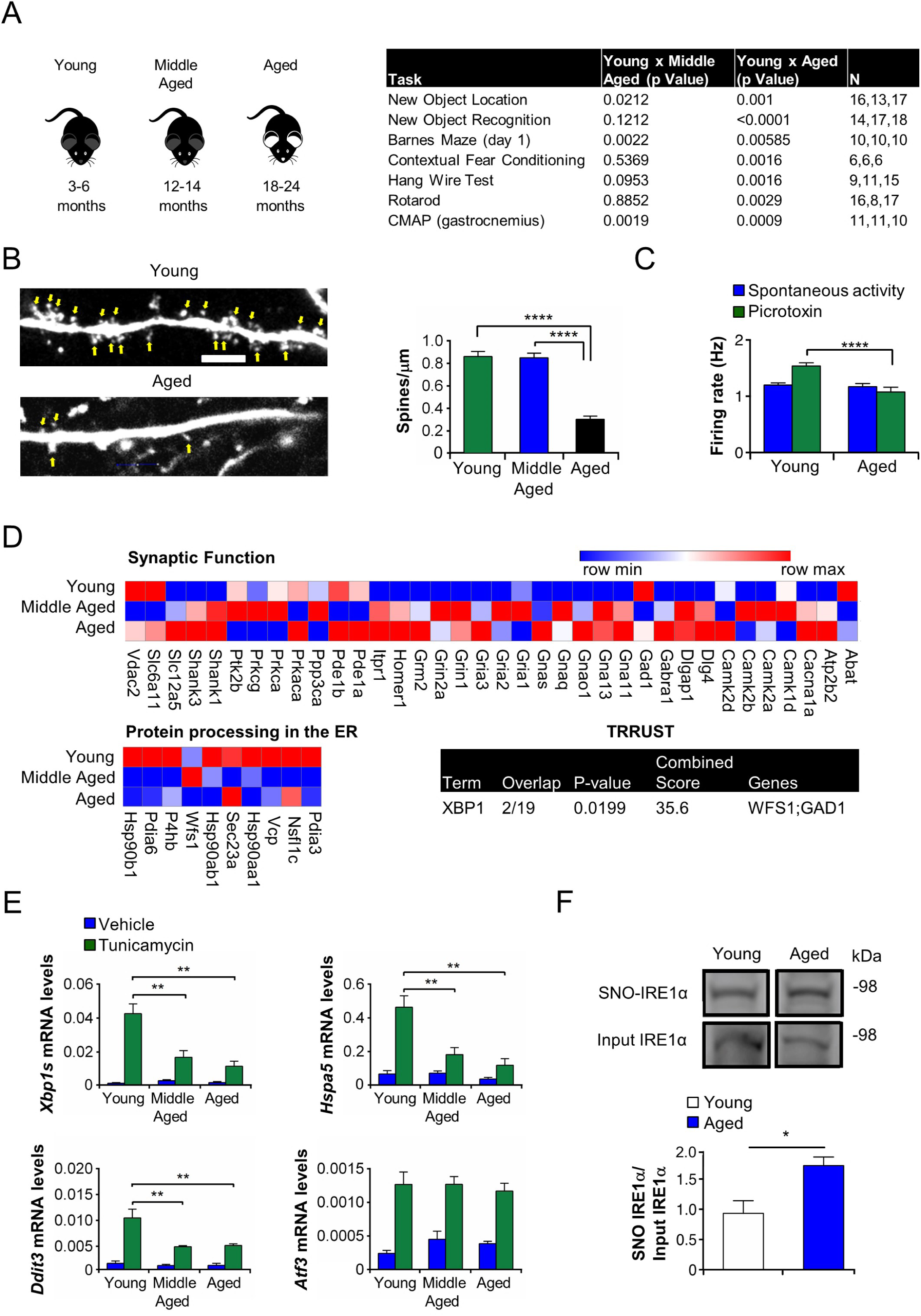
Mapping proteomic and functional alterations during mammalian brain aging. (**A**) Cognitive and motor tasks were performed to monitor age-associated decline in mice. Young (3-6 month-old), middle-aged (12-14 month-old) and aged (18-24 month-old) animals were assessed in the new object location test (NOL), new object recognition test (NOR), contextual fear conditioning (CFC), Barnes maze, wire hanging test, and rotarod performance. Electrophysiological measurements monitored compound muscle action potentials (CMAPs) in gastrocnemius, tibialis anterior, and triceps muscles. Table lists *P* values and number of animals evaluated per group in each functional test. One-way ANOVA followed by Tukey’s post-test was performed as a test of statistical significance. (**B**) Animals of various ages were injected with adeno-associated vector (AAV) into the CA1 region of the hippocampi bilaterally to express eGFP in neurons. One month later, dendritic spines (indicated by arrows) were imaged by confocal microscopy. 60x magnification; scale bar, 5 μm. Left panel: representative fluorescent images showing dendritic spines of pyramidal neurons in CA1 region of AAV-eGFP-injected animals (labeled AAV-Mock). Right panel: Mean and SEM of spine density per μm (*n* = 3 animals/group, *n* = 6-16 dendrites/animal). One-way ANOVA followed by Tukey’s post-test was performed. \*\*\*\**P* < 0.001. (**C**) Brain slice electrophysiological analysis assessing firing rates in CA1 pyramidal neurons (*n* = 4 animals/group; *n* = 832-1584 neurons/animal). Mean and SEM of firing rates were measured during spontaneous activity or following picrotoxin treatment. Unpaired Student t test compared picrotoxin- treated groups. \*\*\*\**P* < 0.001. (**D**) Highly significant proteomic changes comparing the hippocampus of young to middle-aged animals using a cut-off of *P* < 0.05, *n* = 4 animals/group. Functional enrichment analysis on KEGG database using EnrichR platform. Genes enriched in KEGG pathway related to “synaptic function” were grouped in one heatmap (up), and genes enriched in KEGG pathway “protein processing in the ER” were grouped in a separate heatmap (bottom) showing average protein expression in young, middle-aged, and aged samples (*n* = 3-4 animals/group). Bottom right: Prediction of possible transcriptional regulatory interactions inferred from proteomic alterations of young and middle-aged mice using TRRUST version 2 software for both human and mouse databases. Overlap of genes, *P* values, combined scores, and candidate genes of XBP1 were generated by EnrichR platform. (**E**) Young, middle-aged, and aged animals were treated with tunicamcyin (5 mg/kg) or vehicle for 24 h. Following euthanasia, their hippocampi were dissected for RNA extraction. Quantitative PCR showing *Xbp1-s, Hspa5/Bip, Ddit3/Chop* and *Atf3* relative mRNA levels of treated animals compared to controls (*n* = 4 animals/group. One-way ANOVA followed by Tukey’s post-test compared tunicamycin treated groups, \*\**P* < 0.01). **(F)** Representative levels of S- nitrosylated IRE1α and total input IRE1α measured by biotin-switch assay in brains from young and aged mice. Histogram shows relative levels of SNO-IRE1α/input IRE1α for young (3 month-old) vs. aged (18-24 month-old) mouse brains (*n* = 6, \**P* < 0.05 by unpaired Student’s t test).

Since most of the cognitive tasks are dependent on normal hippocampal function, we evaluated the distribution of dendritic spines in CA1 pyramidal cells because their density correlates with the learning capacity *(7).* We found that aged mice manifested significantly lower spine density (Fig. 1B). In agreement with these results, analysis of firing rates of CA1 neurons indicated a decline in the electrophysiological activity of aged mice following picrotoxin (PTX) treatment, an antagonist of GABAergic inhibitory interneurons that fosters excitatory activity in hippocampal circuits (Fig. 1C).

To study molecular alterations in the aging hippocampus, we performed quantitative proteomics to compare tissues derived from animals at different ages. Consistent with the functional decay observed in middle-aged mice, pathway enrichment analysis of significant hits (cut-off: *P* < 0.05) comparing young to middle-aged samples in the KEGG database indicated that neuronal plasticity-related proteins were the most affected (Fig. 1D). Altered pathways included glutamatergic and GABAergic synapses, long-term potentiation (LTP) and calcium signaling (Fig. S2A, tables S1 and S2). Interestingly, enrichment analysis also highlighted proteostasis-related terms such as protein processing in the endoplasmic reticulum (ER) and endocytosis (Fig. S2A and table S2). Analysis of proteomic data with an unbiased text-mining approach using the TRRUST v.2 database *(8)* highlighted the possible contribution of the transcription factor X-Box binding protein 1 (XBP1) in the control of age-related gene expression changes (Fig. 1D, table S2, and fig. S2B). Additionally, XBP1 downstream targets related to ER function and neuronal physiology were also shown to be significantly altered comparing young and middle-aged samples (*9,10*) (Fig. 1D, fig. S2B, and table S2).

XBP1 is a central mediator of the unfolded protein response (UPR), an adaptive pathway that mediates proteostatic recovery in cells suffering ER stress *(11)*. XBP1 is activated by the stress sensor inositol-requiring enzyme-1 alpha (IRE1), an ER-located RNase that catalyzes the unconventional splicing of the XBP1 mRNA to eliminate a 26- nucleotide intron (*11,12*). This processing event shifts the coding reading frame to generate an active and stable transcription factor termed XBP1s (for the <u>s</u>pliced form) *(12)*. XBP1 controls the expression of various components of the proteostasis network *(9)*, but has been also linked to the regulation of synaptic plasticity (*13*). Importantly, studies in simple model organisms suggested that the activity of the UPR declines with aging and that enhanced neuronal XBP1s expression extends health and lifespan in *C. elegans* (reviewed in (*14*)).

To empirically test the capacity of the aging brain to engage the IRE1α-XBP1s pathway, we intraperitoneally injected animals of different ages with tunicamycin, a well-established inducer of ER stress *(15)* (Fig. 1E). Next, we evaluated both IRE1α and general UPR transcriptional responses by measuring the mRNA levels of *Xbp1s, Hspa5* (*Bip/Grp78*), *Ddit3* (*Chop*) and *Atf3*. Remarkably, the capacity to induce *Xbp1*s was reduced in hippocampus in middle and old age following the induction of ER stress (Fig. 1E). UPR mediators *Ddit3* and *Hspa3* showed the same trend in this brain region (Fig. 1E). However, these findings were not recapitulated when the cerebellum of the same animals was analyzed, whereas the cerebrocortex showed mild but significant effects (fig. S2,C and D). These findings suggest that the occurrence of specific molecular alterations in the aging hippocampus interfere with the capacity to adapt to ER stress. Along these lines, increased generation of reactive oxygen and nitrogen species, such as nitric oxide (NO), has been observed during aging (*16*). Previous reports indicated that S-nitrosylation of IRE1α, resulting from posttranslational modification of cysteine thiol groups at Cys931 and Cys951 by NO-related species, inhibits its ribonuclease activity (*17)*. Accordingly, we evaluated the levels of S-nitrosylation of IRE1α (abbreviated SNO-IRE1α) during normal brain aging. We measured the ratio of SNO-IRE1α to total IRE1α in the brains of young and old animals, and found that the ratio significantly increased in aged samples (Fig. 1F), potentially accounting, at least in part, for the decrease in IRE1α activity and thus compromised ability to adapt to ER stress with age.

To assess the significance of the UPR to brain health span, we conditionally ablated the RNase domain of IRE1α in the nervous system using CRE transgenic lines driven by the Nestin promoter (IRE1^cKO^) for general deletion in the brain *(18)* or the Camk2a promoter (IRE1^cKO/CaK^) (Fig. S3A) to restrict the targeting to specific neuronal populations. Disruption of the IRE1α pathway in the brain resulted in reduced performance in age-sensitive cognitive tests, including new object recognition (NOR) and contextual fear conditioning (CFC) (Fig. 2, A and B).

**Figure 2.**
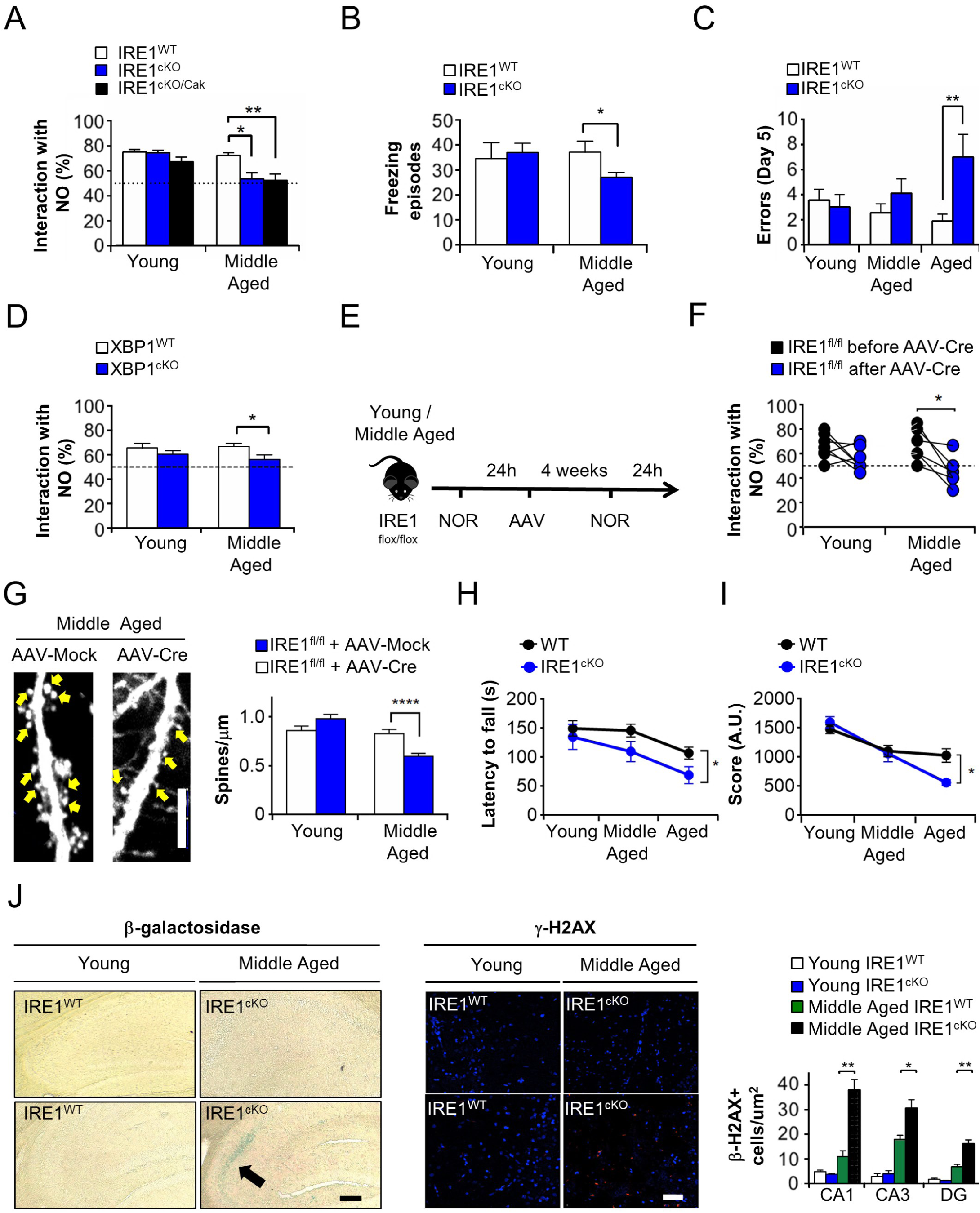
Genetic ablation of the IRE1a pathway in the brain accelerates and exacerbates age-associated cognitive and motor decline in mammals. **(A)** Conditional knockout mice for IRE1α were generated using the Nestin-CRE system (IRE1^cKO^), their littermate flox/flox control animals (IRE1^WT^), or a second conditional knockout animal using calcium/calmodulin-dependent protein *kinase* II alpha-CRE (IRE1^cKO/Cak2^). The new object recognition test was used to evaluate the ability to discriminate novel objects after 24 h in young (*n* = 10, 10, 8) and middle-aged mice (*n* = 10, 10, 10; \**P* < 0.05, \*\**P* < 0.01 by one way ANOVA followed by Tukey’s post-hoc test comparing each age matched group). (**B**) In animals of indicated genotypes, contextual fear conditioning test was performed to compare total number of freezing episodes following 24 h of aversive stimuli in young (*n* = 7, 10) and middle-aged (*n* = 9, 14) wild-type (IRE1^WT^) or IRE1^CkO^ mice (\**P* < 0.05 by unpaired Student’s t test within each age-matched group). (**C**) Young (*n* = 9, 9), middle-aged (*n* = 9, 9), and aged (*n* = 10, 8) IRE1^WT^ or IRE1^cKO^ mice were evaluated in the Barnes maze to evaluate spatial memory acquisition. Total number of errors before finding the target hole is plotted (\**P* < 0.05 by unpaired Student’s t test within each age-matched group). **(D)** Young WT (*n* = 8), young XBP1s^cKO^ (*n* = 12), middle-aged WT (*n* = 18), or middle-aged XBP1^cKO^ (*n* = 14) mice were evaluated in the new object recognition test to analyze the ability to discriminate between novel objects following 24 h of presentation of two identical objects (\**P* < 0.05 by unpaired Student’s t test comparing age-matched groups). (**E**) Scheme to illustrate the loss-of-function approach based on AAV2-CRE-mediated deletion of floxed IRE1α in the hippocampus. IRE1^flox/flox^ animals were evaluated using the NOR test and then received bilateral hippocampal injections with AAV2-CRE. After 4 weeks, animals were evaluated again in the NOR test and then euthanized. Brains were collected for dendritic spine and biochemical analysis. (**F**) Young (*n* = 7) and middle-aged (*n* = 6) IRE1^flox/flox^ animals were evaluated in the NOR test before and after AAV2-CRE intra-hippocampal injections (\**P* < 0.05 by paired Student’s t test comparing each individual before and after AAV2 injection). (**G**) Animals of different ages were injected with adeno-associated vector (AAV-CRE or AAV2-Mock) into the CA1 region of each hippocampus. AAV constructs included a GFP cassette to express eGFP in neurons for monitoring dendritic spine density. One month after the injection, spines were imaged by confocal microscopy. Left-hand panels: Representative confocal microscopic images of dendritic spines (arrows) in the CA1 region of middle-aged animals. 60x magnification; scale bar, 5 μm. Right-hand panel: Histogram of mean and SEM of spine density per μm (*n* = 24 dendrites, 4 animals; 27 dendrites, 3 animals; 31 dendrites, 4 animals; 42 dendrites, 4 animals, respectively). \*\*\*\**P* < 0.001 by unpaired Student’s t test within each age-matched group. (**H**) Wire hanging test performed in young, middle-aged, and aged IRE1^WT^ or IRE1^cKO^ to evaluate motor performance. Graph indicates mean and SEM of arbitrary scores from aged (*n* = 18, 15), middle-aged (*n* = 11, 15), and young (*n* = 10, 12) animals (\**P* < 0.05 by two-way ANOVA followed by Sidak’s post-hoc test). (**I**) Rotarod test used to evaluate motor performance and coordination in young (*n* = 16, 12), middle- aged (*n* = 8, 8), and aged (*n* = 18, 6) IRE1^WT^ or IRE1^cKO^ animals. Graph indicates mean and SEM. of latencies to fall from the rod (\**P* < 0.05 by unpaired Student’s t test within each age group). **(J)** Representative photomicrographs of β-galactosidase staining (at left, indicated by arrow) and immunofluorescence for γ-H2AX (middle panel, red staining, representing a biomarker for DNA damage), counterstained for cell nuclei (blue) of hippocampal slices derived from young and middle-aged IRE1^WT^ or IRE1^cKO^ animals (*n* = 3-4 animals/group). Graphs (at right) indicate mean and SEM of percentage of γ-H2AX- positive cells. \**P* < 0.05, \*\**P* < 0.01 by unpaired Student’s t test within each age-matched group for each hippocampal sub-region. β-galactosidase magnification, 20x; scale bar, 200 μm. Immunofluorescence for γ-H2AX, magnification, 40x; scale bar, 50 μm.

Importantly, aged mice interacted with objects for the same period of time as young animals (fig. S1B and 1F), and did not show intrinsic preferences for any of the objects or locations tested (fig. S1B-G), excluding the possibility of age-associated motor dysfunction. Remarkably, only aged IRE1^cKO^ animals presented a significant decay in spatial memory acquisition when tested in the Barnes maze, reflected in a higher percentage of errors (Fig. 2C), although the latency to find the targets did not differ between genotypes (fig. S3B). Importantly, genetic disruption of IRE1α function did not alter the cognitive performance of young animals, indicating the occurrence of age-dependent phenotypes (Fig. 2A-C, Fig. S3, B and C). Since IRE1α has various signaling outputs in addition to controlling XBP1 mRNA splicing (*19*), we further confirmed our results, finding that cognitive performance with the NOR assay in XBP1 conditional knockout animals, which have normal IRE1α expression but lack XBP1 expression in the hippocampus, was also impaired starting in middle age but not in young animals (Fig. 2D).

We next targeted the UPR in the brain of adult animals via local delivery of CRE recombinase into the hippocampus of IRE1α floxed animals using adeno-associated viruses (AAVs) with serotype 2 (Fig. 2E and fig. S3D). Middle-aged mice were tested in the NOR assay prior to AAV-CRE injection and then monitored again following 4-weeks of brain surgery. Remarkably, targeting IRE1α in the hippocampus of middle-aged mice impaired the capacity to discriminate novel objects (Fig. 2F), correlating with reduced density of dendritic spines in the CA1 region (Fig. 2G). Importantly, injection of AAV- CRE into the brain of young IRE1^flox/flox^ animals did not alter NOR performance or the distribution of dendritic spines (Fig. 2, F and G), confirming the occurrence of age-related impairment.

We then determined whether IRE1α deficiency in the brain exacerbates decay in motor ability and coordination during aging. Aged IRE1^cKO^ mice manifested impaired performance in the rotarod and wire hanging tests when compared to littermate control animals (Fig. 2, H and I). However, muscle electrophysiological properties, assessed by compound muscle action potential (CMAP) amplitudes, did not show significant differences between genotypes in three distinct muscles tested (fig. S3E-G). Similarly, analysis of neuromuscular junction (NMJ) morphology did not reveal any alterations between aged IRE1^cKO^ and control mice (fig. S3, H and J). Taken together, these results suggest that motor defects triggered by IRE1α deficiency are not mediated by postsynaptic muscular innervation but rather by presynaptic mechanisms. Finally, we evaluated the accumulation of senescent cells, as indicated by DNA damage, in hippocampal tissue of IRE1^cKO^ mice as a measure of a biological marker of aging. Remarkably, an increase in senescent cells was observed in the brain of middle-aged IRE1^cKO^ animals but not in young animals (Fig. 2J and fig. S3K). Overall, our results indicate that genetic disruption of the IRE1α pathway in the nervous system accelerates the natural emergence of age-associated behavioral and neuromorphological alterations.

In order to test the consequences of improving neuronal proteostasis, we exogenously bolstered an adaptive UPR by expressing the spliced and active form of XBP1 in the brain. For this purpose, we initially used transgenic mice that overexpress XBP1s under the control of the PrP promoter *(13*) (referred to here as Tg^XBP1s^; fig. S4A) and evaluated their cognitive performance during aging. Remarkably, XBP1s overexpression prevented the development of age-related deterioration in brain function, as evaluated in the NOR, NOL and Barnes maze tests; in fact, these mice performed comparably to non-transgenic young animals (Fig. 3A-C). Additionally, Tg^XBP1s^ mice showed reduced age-dependent coordination and motor decay when compared to littermate controls in the wire hanging (Fig. 3D) and rotarod tests (Fig. 3E).

**Figure 3.**
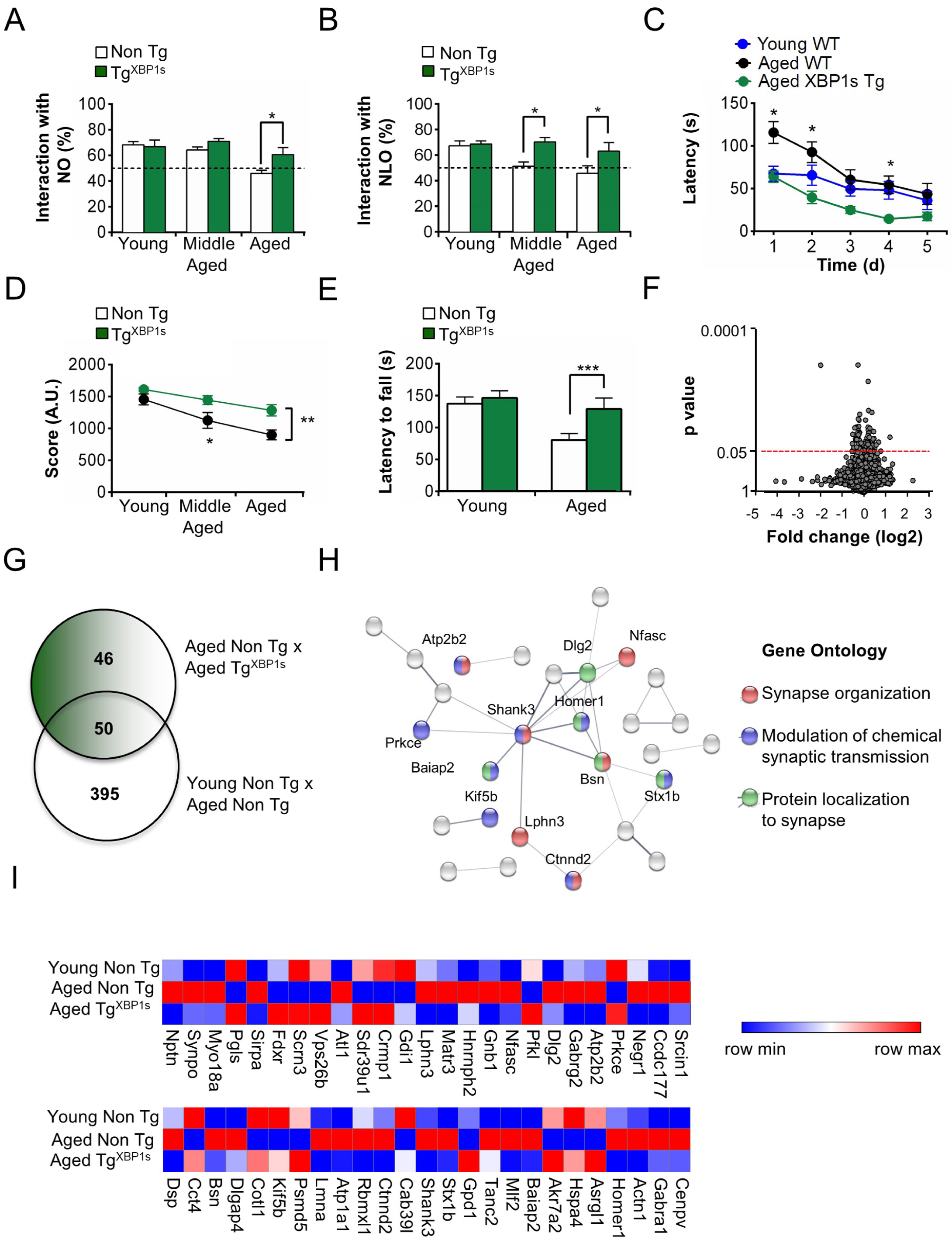
XBP1s overexpression in the brain prevents age-associated motor and cognitive decline in mammals. (**A**) New object recognition test was used to evaluate ability to discriminate novel objects after 24 h in Tg^XBP1s^ or littermate non-transgenic animals (non-Tg) in young, middle-aged, and aged animals (*n* = 22, 6; 23, 9; 17, 12; respectively; \**P* < 0.05 by unpaired Student’s t test within each age-matched group). (**B**) New object location test was used to evaluate the ability to discriminate changes in object location after 24 h in young, middle-aged, and aged non-Tg or Tg^XBP1s^ (n = 20, 7; 12, 13; 19, 9 respectively; histogram shows mean and SEM of percentage of time interacting with the novel-located object (NLO). \**P* < 0.05 by unpaired Student’s t test within each age-matched group). (**C**) Aged non-Tg and aged Tg^XBP1s^ animals were evaluated in the Barnes maze test to compare spatial memory acquisition over a period of 5 days. Graph indicates mean and SEM of each group (*n* = 10 for non-Tg [WT], 8 for Tg^XBP1s^). Student’s t test was used to compare performances within each day (\**P* < 0.05). Latencies of young/non-Tg mice also plotted for comparison (*n* = 10). (**D**) Young, middle-aged, and aged non-Tg or Tg^XBP1s^ were evaluated in the wire hanging test to monitor motor performance and coordination. Mean and SEM of arbitrary scores for each group (*n* = 8 animals/group; \**P* < 0.05, \*\**P* < 0.01 by two-way ANOVA followed by Sidak’s multiple comparison test). **(E)** Young and aged non-Tg or Tg^XBP1s^ animals were evaluated in the rotarod test to monitor motor performance and coordination. Mean and SEM for each group (*n* = 10 animals/group, \*\*\**P* < 0.001 by unpaired Student’s t test within each matched group). (**F**) Volcano plot of proteomic alterations in the hippocampus of aged non-Tg and aged Tg^XBP1s^ animals indicates *P* values (y-axis) and fold change (log2, x- axis). Red dotted lines delineate cut-off used to filter genes for functional enrichment analysis. (**G**) Venn diagram showing most significant proteomic alterations (cut-off: *P* < 0.05) between comparisons of young non-Tg, aged non-Tg, and aged Tg^XBP1s^ animals. (**H**) Protein-protein interaction networks generated by STRING v.11 (left) using most significant proteomic changes comparing the hippocampus of aged non-Tg with aged Tg^XBP1s^ (cut-off: *P* < 0.05, *n* = 4 animals/group). Network edge line thickness indicates strength of data support. Only connected nodes are shown. Enrichment analysis in Gene Ontology database was performed, and most enriched terms are indicated (right). Proteins related to each enriched term are colored in red, blue or green in the network (number of nodes: 50, number of edges: 32, average node degree: 1.28, average local clustering coefficient: 0.427, PPI enrichment *P*-value: 1.48e-05). (**I**) Heat maps indicate significant gene alterations comparing young non-Tg, aged non-Tg, and aged Tg^XBP1s^. Most significant proteomic changes, comparing the hippocampus of aged animals, were determined using a cut-off of *P* < 0.05 (*n* = 4 animals/group).

We then analyzed global proteomic changes in hippocampal tissue derived from aged (Fig. 3, F and G) and middle-aged Tg^XBP1s^ animals (fig. S4, B and C) versus age-matched controls. Enrichment analysis and protein-protein interaction network assessment revealed that overexpression of XBP1s in the aging brain altered the expression of proteins related to synapse organization, modulation of chemical synaptic transmission, and protein localization to synapses (Fig. 3H, tables S3 and S4). Consistent with our behavioral data, overexpression of XBP1s in the brain of aged (Fig. 3I) and middle-aged (fig. S4D) animals prevented the occurrence of most age-related proteomic alterations. To validate the proteomic data, we measured the levels of GAD1 by western blot and confirmed its predicted expression in hippocampal tissue of Tg^XBP1s^ during aging (fig. S4E). Our results are consistent with the notion that XBP1s overexpression prevents brain proteome alterations, thus preserving normal synaptic function during aging.

We next evaluated whether artificial activation of XBP1s-dependent responses could potentially reverse the natural decay in normal brain function during aging. For this purpose, we performed bilateral injections of AAVs to express XBP1s in the hippocampi of aging mice that already manifested cognitive decline (fig. S4, F and G). Remarkably, the administration of AAV-XBP1s to these middle-aged and aged mice resulted in improved performance in the various cognitive tests compared to age-matched animals injected with control virus (Fig. 4A-D).

**Figure 4:**
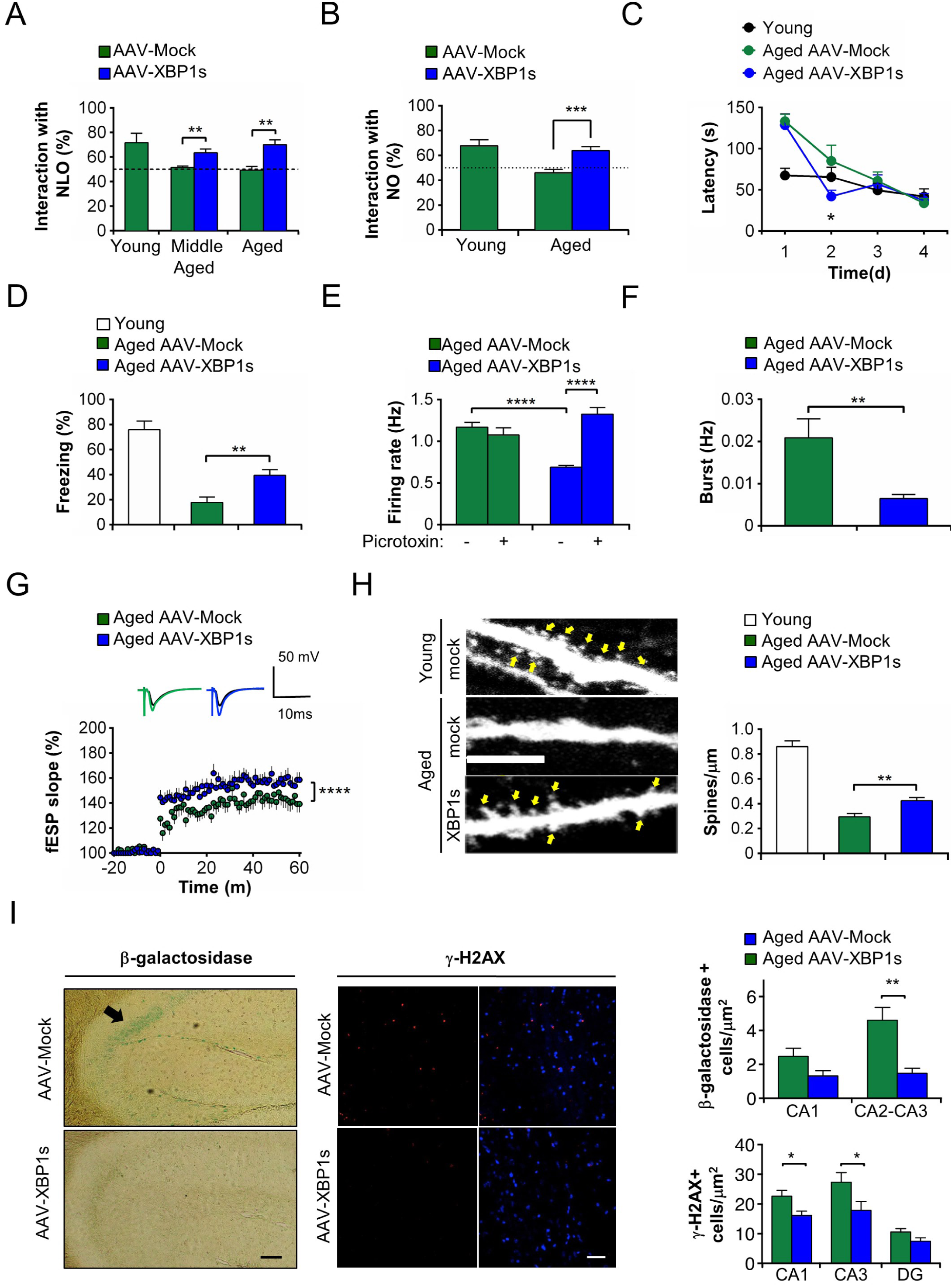
XBP1s gene delivery to middle-aged or aged WT animals reverses age-associated phenotypes at behavioral, morphological and electrophysiological levels. (**A**) Middle-aged and aged WT animals were injected with AAV2-Mock or AAV2-XBP1s in both hippocampi and then evaluated in the new object location test to evaluate their ability to discriminate displacement of objects 24 h following presentation. Histogram shows mean and SEM of percentage of time interacting with the novel located object (NLO) for *n* = 10 animals/group, \*\**P* < 0.01 by unpaired Student’s t test). Young animal performance plotted for comparison (*n* = 6). (**B**) Aged animals injected with AAV2- Mock or AAV2-XBP1s in both hippocampi were evaluated in the new object recognition test to evaluate their ability to discriminate novel objects after 24 h. Mean and SEM of percentage of interaction time with novel objects (*n* = 10 animals/group, \*\*\**P* < 0.005 by unpaired Student’s t test). Young animal performance plotted for comparison (*n* = 6). **(C)** Aged animals injected with AAV2-Mock or AAV2-XBP1s in both hippocampi were evaluated in the Barnes maze test to compare spatial memory acquisition. Mean and SEM of latencies to find target for each group during 4 days of testing (*n* = 5 for AAV2-Mock, 8 for AAV2-XBP1s; Student’s t test was used to compare performances within each day (\**P* < 0.05). Young animal performance plotted for comparison (*n* = 10). **(D)** Aged animals injected with AAV2-Mock or AAV2-XBP1s evaluated by contextual fear conditioning for aversive memory acquisition 24 h after presentation of an unconditioned stimulus (*n* = 5 for AAV2-Mock, 7 for AAV2-XBP1s; \*\**P* < 0.01 by unpaired Student’s t test). Young animal performance plotted for comparison (*n* = 6). **(E)** Brain slice electrophysiological analysis assessing firing rates in hippocampal neurons. Mean and SEM of firing rates were measured during spontaneous activity or following picrotoxin treatment in brain slices from aged mice injected with AAV2-Mock or AAV2-XBP1s (*n* = 633-832 neurons; *n* = 6, 9 animals, respectively; \*\*\*\**P* < 0.001 by unpaired Student’s t test). **(F)** Burst activity measured in pyramidal and interneurons in hippocampal brain slices derived from aged mice injected with AAV2-Mock or AAV2-XBP1s (*n* = 832, 644 neurons from 3-4 animals, \*\**P* < 0.01 by unpaired Student’s t test). **(G)** Mean and SEM of field excitatory postsynaptic potentials in brain slices derived from aged animals injected with AAV2-Mock or AAV2-XBP1s 1 h after theta burst stimulus (TBS) to induce LTP in (*n* = 6, 9 animals, respectively; *n* = 17-28 slices/animal, \*\*\*\**P* < 0.001 by unpaired Student’s t test). **(H)** Aged mice were injected with adeno-associated vector (AAV) into CA1 region hippocampus to express eGFP. One month later, dendritic spines (arrows) were imaged by confocal microscopy (60x magnification; scale bar, 5 μm). Left panel: Representative fluorescent images showing dendritic spines of pyramidal neurons in CA1 region of young and aged animals injected with AAV2-eGFP (mock) or aged animals injected with AAV2-XBP1s. Right panel: Mean and SEM of spine density per μm for indicated experimental groups (*n* = 24, 53 and 35 dendrites from 3, 6 and 5 animals, respectively; \*\**P* < 0.01 by unpaired Student’s t test comparing the aged groups). Young animal dendritic spine per length plotted for reference. 60x magnification; scale bar, 5 μm). (I) Representative images of β-galactosidase staining (left) and immunofluorescence for γ-H2AX (middle) of hippocampal slices derived from aged mice injected with AAV2 (*n* = 3-4 animals/group). Histograms (right) show mean and SEM of percentage of γ-H2A- positive cells. β-galactosidase magnification 20x;scale bar, 200 μm. Immunofluorescence for γ-H2AX magnification, 40x; scale bar, 50 μm. \**P* < 0.05, \*\**P* < 0.01 by unpaired Student’s t test comparing the aged groups.

The beneficial effects of AAV-XBP1s treatment correlated with the improved electrophysiological properties of hippocampal slices (Fig. 4, E and F). In fact, we observed that basal firing rates and bursting activity in hippocampal neurons were decreased after treating aged animals with AAV-XBP1s. Such alterations facilitated increased firing rates in CA1 neurons following PTX treatment (Fig. 4, E and F). Furthermore, induction of LTP, widely thought to represent an electrical correlate of learning and memory, was significantly improved in hippocampal slices derived from aged mice treated with AAV-XBP1s compared to controls (Fig. 4G and fig. S4H). Importantly, these findings were associated with a significant increase in dendritic spine density in CA1 neurons compared to age-matched control animals (Fig. 4H). Interestingly, we also found that aged mice presented decreased accumulation of senescent cells in the hippocampus following XBP1s overexpression (Fig. 4I). Overall, our results indicate that XBP1s gene transfer to the aged brain is sufficient to rescue age-associated cognitive dysfunction at behavioral, electrophysiological and morphological levels.

Prior functional studies have demonstrated the involvement of the IRE1α-XBP1s pathway in a variety of age-related neurodegenerative conditions, including Parkinson’s disease, Alzheimer’s disease, and frontotemporal dementia (reviewed in *19*). Studies in invertebrate models (yeast, *C. elegans* and *D. melanogaster*) have uncovered a central role of ER proteostasis and the UPR in aging *(20-25)*. Moreover, the beneficial effects of caloric restriction in extending life and health span in simple model organisms have been recently linked to modulatory effects on ER proteostasis (*26-28*). Heretofore, however, only correlative studies have associated aging and ER stress for mammals, albeit in multiple organs and tissues (*14*), and a causal link has remained speculative, although XBP1 deficiency was shown to accelerate retinal degeneration in diabetic mice during aging (*29*). Here, we demonstrate for the first time a fundamental role for the IRE1α- XBP1s pathway in maintaining brain function during normal aging. XBP1s overexpression prevented age-mediated proteomic alterations, including crucial components of glutamatergic and GABAergic synapses (*30*). One of the altered proteins, GAD-1 (glutamate decarboxylase), converts glutamate into GABA and is a downstream target of XBP1 in human tissue (*31*). Surprisingly, overexpression of XBP1s in the hippocampus did not result in clear upregulation of canonical UPR target genes described in other tissues. This is in agreement with recent findings suggesting that the UPR, and more specifically XBP1, has alternative functions in the nervous system, directly controlling synaptic plasticity (*32*). Importantly, the appearance of senescent cells, which is associated with age-mediated loss of brain function (*33*), was prevented by XBP1s overexpression, whereas IRE1α deficiency accelerated the accumulation of senescent cells, suggesting that the UPR exerts global effects in sustaining the health of brain tissue during aging.

Recently, the neuronal UPR has been suggested as a central regulator of organismal proteostasis in simple model organisms through a cell-nonautonomous mechanism that tunes proteostasis in the periphery (*21-23,25*). It remains to be determined, however, whether an increase in XBP1s expression in the aged brain can be translated into the propagation of adaptive signals that improve the function of other organs, thus mitigating their deterioration during the course of natural aging. In conclusion, our results indicate that intervention strategies that improve ER proteostasis via the UPR may extend brain health span, reducing the risk of developing dementia and age-associated neurodegeneration in the elderly.

## Acknowledgments

We thank Dr. Takao Iwawaki for providing IRE1α null animals. We thank Javiera Ponce, Francisco Aburto and Diego Rivas for animal care. We also thank Monica Flores and Maria Painavel for technical support.

## Author contributions

FCM, CH: **Conceptualization, Formal analysis, Funding acquisition, Investigation, Methodology, Software, Experimentation, Project administration, Visualization, Writing – original draft**. GT, GM, DM, YG, CDA, TM, AOA, CG, CS, FBG, SA, KV, HH, XZ, TN: **Investigation, Methodology, Formal analysis, Project administration, Visualization, Data curation**. FCM, GM, DM, YG, CDA, SPS, BK, JCC, AGA, LP, JPH, CH, SPS, BK, JCC, AP, LP, JPH, SAL: Funding Acquisition, **Resources, Formal analysis, Investigation, Methodology, Supervision, Writing - review & editing.**

## Data and materials availability

All data are available in the main text or the supplementary materials.

