## Supplementary Material for "Control of mammalian brain aging by the unfolded protein response (UPR)"

**Affiliations:**

**This file includes:**

Materials and Methods

References for Materials an Methods

Figs. S1 to S4

Tables S1 to S4

**Materials and Methods**

**Animals**

C57BL/6 mice were employed, maintained in a facility with 12 h light/dark cycle at 25°C with food and water provided ad libitum. Cohorts of aged, middle-aged, and young animals were directly obtained from the Jackson´s Laboratory (USA). All animal procedures were approved by the Bioethics Committee of the Faculty of Medicine, University of Chile (protocol number 18166-MED-UCH).

**Stereotaxic injections**

Young (3 month-old), middle-aged (12 month-old), and aged (18 month-old) mice received bilateral stereotaxic injections of AAV (1 × 10^9^ viral genomes (VGs)/μL) in the hippocampi using the coordinates AP: -1.9, DV: +1.7, and ML: ±1.0. Male mice were deeply anesthetized using isoflurane (4%) and a stereotaxic apparatus coupled to a Hamilton microsyringe was used for the procedure. Cranium was exposed though a skin incision, and bilaterally symmetrical holes were opened using a dental drill. Injections were performed at approximately 0.5 μL/min in a total of 1 l. Mice were returned to their home cages and kept under close monitoring until they were awake. Mice were injected with either control AAV serotype 2 vector (Mock), AAV2-CRE (Addgene, #105545-AAV2) or AAV2-XBP1s (produced at Genzyme) under the control of the CMV promoter, in addition to an eGFP cassette to monitor transduction efficiency, as we previously reported (*1*).

**Behavioral tests**

Behavioral experiments were performed in a masked fashion, both for genetically-modified animals and AAV-injected mice, using groups of age-matched controls. Injected mice underwent behavioral evaluation 1 month following injections. Cognitive and motor tests were performed in the following order: New object recognition, new object location, Barnes maze, wire hanging test, rotarod, and contextual fear conditioning. Not all animals were exposed to all tests in order to avoid possible secondary effects mediated by interaction of the various tests.

1. *Novel object recognition*

Novel object recognition (NOR) was performed as previously described *(1-2).* Briefly, mice were first trained and then placed in an arena facing the wall. They were presented with two identical objects (Lego blocks, 3 x 4 cm) located in a distal position towards the animal. For the training phase, mice were allowed to explore both objects for a total time of exploration of 20 s during a single trial. If exploration did not occur within 20 s, a maximum time of 5 minutes was given for each animal to explore the objects. After 24 h, mice were again placed in the arena facing the wall but at this time were introduced to a new object of different color, shape and texture. In the test phase, mice were again allowed to explore objects for a maximum of 20 s or 5 minutes if the criterion of exploration was not accomplished in the initial time allotment. Four different Lego blocks with equivalent sizes (3-4 x 4 x 4 cm) were presented in a randomized fashion for each animal. Trials were recorded on a digital camera coupled to a computer for subsequent evaluation of time spent exploring objects by a blinded researcher. Both the objects and the arena were cleaned with 70% ethanol before each session to avoid olfactory cues. Exploration time was defined as the amount of time mice had their noises oriented toward an object with its nose within 3 cm or less. Other behaviors such as rearing near the object or resting against the object were not considered as exploration. Exploration time spent during the training phase was also recorded. Animals that failed to interact with one or both objects, or that showed signs of stress were excluded from the analysis. All analysis were performed in a masked fashion.

2. *Novel object location*

The novel object location task (NOL) was performed as previously described *(2-3).* Mice were placed in the same arena used for the NOR assay. The arena was divided into quadrants. For the training phase, mice were introduced to two identical objects (Lego blocks, 5 x 4 x 4 cm) placed in two randomized quadrants of the arena. Objects were placed equidistant from one other. At 3 or 24 h after the test phase, mice were again placed in the arena facing the wall but now with one of the objects placed in a distinct quadrant. Trials were recorded on a digital camera coupled to a computer for subsequent evaluation of time spent exploring objects. Both the objects and the arena were cleaned with 70% ethanol before each session to avoid olfactory cues. Exploration time was defined as detailed above by a blinded researcher. As per standard protocol, animals that failed to interact with one or both objects or showed signals of stress were excluded from the analysis.

3. *Barnes maze*

The Barnes maze task was performed on a white circular surface (0.9 m in diameter) with 20 holes equally spaced around the perimeter, as previously described *(4-5).* A dark escape box (10 × 20 × 7.8 cm) was located under one of the holes, which was the target. A ramp was placed under the target hole so that mice could reach the escape tunnel easily. The circular open field was elevated 75 cm above the floor. Distal spatial cues (with different colors and shapes) were placed outside of the maze. The maze was rotated daily, with the spatial location of the target unchanged with respect to the distal visual room cues. A cylindrical start chamber was placed in the center of the maze and removed after 10 s. Training sessions consisted of four trials per day conducted for 4 days with a maximum time of 3 minutes each. If animals were unable to find the target after 3 minutes, they were gently placed in the right hole by the tail. A stopwatch sound was used as an aversive stimulus in order to induce exploration. Once a mouse reached the target, the noise was immediately stopped, and the mouse was left inside the box for 1 minute and then returned to its home cage. A maximal period of 15 minutes was given for the inter-trial interval. The maze and the escape box were cleaned with 70% ethanol between each trial to avoid olfactory cues. The numbers of primary pokes and the primary latency to reach the target were manually counted, as defined by the number of pokes and time to reach the target for the first time, as some animals did not promptly enter the escape box during training sessions (*4*). Every trial was recorded on a video camera placed in the ceiling and fed to a computer. On the fifth day, the escape box was removed, and one single trial was performed as a probe trial to evaluate spatial memory acquisition with a maximum time of 90 s. Then, following 7 days without training, mice were again exposed to a test phase to evaluate long-term memory formation.

4. *Contextual fear conditioning*

On the first day, mice were placed in the contextual fear-conditioning chamber (Med Associates). During the first 2 min, no stimulus was applied, following which 80 db of white noise (the conditioned stimulus; CS) was generated for 30 s. Two seconds later, mice were exposed to a 0.5 mA electric shock (the unconditioned stimulus) for 2 s. After the shock, mice were let in the cage for 2 more minutes ,and freezing episodes, represented by the percentage of freezing response to total activity, were measured by an automated system. On the next day, mice were again placed in the same chamber and exposed to the same protocol but this time without any electroshock. Freezing events and freezing percentage time were recorded for 5 min using an automated system.

5. *Wire hanging test*

The wire hanging test is a well-established protocol for measuring muscular strength in mice. We generated an arbitrary score to take into consideration the number of times one animal fell from the wire during the course of the test (total of 180 s). Every time an animal fell from the wire, its arbitrary score on the test was decreased by 1 (starting at 10 and decreasing to unity). If an animal fell more than 9 times, the test was stopped. Every time an animal reached one side of the wire, the stopwatch was paused, and the animal was placed in the middle of the wire before the stopwatch was started again. A soft cotton sheet was placed under the wire to avoid an harm to the animal during the falls. The test was scored as follows:

*Final Score = (10 – number of falls)* x *time remaining on the wire (s).*

6. *Rotarod test*

Mice were placed on a rotating cylinder and trained to learn how to walk on it at constant speed. Mice that could not learn the task after successive trials were excluded. Mice that learned the task were then analyzed in 5 trials with the apparatus configured to accelerate from 4-40 RPM (rotations per minute) over 10 minutes during the course of 3 days. The mean time until the mouse fell off of the cylinder for each trial was computed as the score.

**Tissue collection and processing**

Briefly, mice were deeply anesthetized with ketamine/xylazine and perfused with ice cold saline. The brain was removed from the skull and separated into two hemispheres. The hippocampus, cerebral cortex and cerebellum of the left hemisphere were dissected out and stored frozen at -80 °C until analysis. The right hemisphere was postfixed in 4% paraformaldehyde in PBS overnight at 4 °C, followed by cryopreservation in 30% sucrose in PBS and freezing medium (OCT, TissueTek). Subsequently, 40 μm-thick sagittal sections were obtained free-floating on a Leica cryostat for immunostaining using anti-NeuN (Millipore, MAB377, 1:100) to label neurons. Five serial sections every 200 μm were stained per animal. Fluorescence images were acquired using a confocal microscope (Nikon C2+)**.**

**Dendritic spine imaging**

Brain slices were cut at 40-m thickness on a cryostat. AAV2-GFP fluorescence was previously confirmed in injected animals to validate viral transduction using an inverted epifluorescence microscope and then imaged on a confocal microscope, Nikon Eclipse T1, at 60x magnification with additional digital zoom of 3x. Similar regions were compared among animals (CA1 region, spines in primary and secondary dendrites between the *stratum radiata* and the pyramidal layer, AP: -1.9 to -2.1 from the bregma). Z-stacks were acquired in 0.5-m slices, laser intensity at 0.5-1% and 12.5us/pixel at 1024x1024 resolution. Z-Stacks were then summed using ImageJ software for total maximum intensity to generate one single stacked 8-bit image. The number of spines was manually quantified in scaled images and divided by the length of the dendrite analyzed; 5-10 dendrites per animal were used for analysis.

**Immunofluorescence**

Brain slices were incubated in citrate buffer at 96 °C for 30 min for epitope exposition and washed in PBS. After this, slices were washed in TBS and incubated in blocking solution (3% BSA and 0.05% Triton X-100) for 60 min at room temperature. Slices were then incubated overnight at 4 °C with primary antibody anti-phospho-Histone H2AX Ser139 (Sigma-Aldrich, 05-636) diluted 1:1000 in 3% BSA in TBS. Secondary antibody was Alexa Fluor 568 (Invitrogen A-11031) diluted 1:2000 in 3% BSA in TBS. The samples were washed 3 times using TBS, and in the final wash, DAPI was added and incubated for 5 minutes. Images were taken with Leica TCS SP8 confocal microscope with a 40X objective magnification. ImageJ and LAS X software were used to process the stacked images. The percentage of positive cells for p-H2AX was graphed. Representative images are shown.

**Senescence-associated beta-galactosidase (SA-βgal) staining**

Histochemical detection of SA-βgal activity was performed according to Debacq-Chainiaux et al. (*6*). Briefly, slices were incubated in a 1 mg/mL of solution of 5-bromo-4-chloro-3-indolyl β-d-galactopyranoside in 0.04 M citric acid/sodium, 0.005 M K_3_FeCN_6_ , 0.005 M K_4_FeCN_6_ , 0.15 M NaCl and 0.002 M MgCl_2_, and diluted in phosphate-buffered saline (pH 6) for 16 h. After incubation, slices were washed with TBS and mounted on superfrost microscope slides (ThermoFisher 6776214) using Fluoromount-G (ThermoFisher, 00-4958-02). Images were acquired on a Leica DM500 binocular microscope equipped with a Leica ICC50 W camera using 4X and 10X objective magnifications. ImageJ software was used to process the images. Positive area for SA-βgal activity was measured and representative images are shown.

**Electrophysiological measurements**

1. *Excitatory postsynaptic field recordings*

Hippocampal slices were prepared as we previously reported *(7, 8)*. Briefly, mice were deeply anesthetized with isoflurane (Forene B506 AbbVie), brains quickly removed, and hippocampi sectioned into 300-μm-thick slices in ice-cold dissection buffer (in mM: 2.6 KCl, 1.23 NaH_2_PO_4_, 26 NaHCO_3_, 212.7 sucrose, 10 dextrose, 3 MgCl_2_, and 1 CaCl_2_, equilibrated with 95% O2 and 5% CO2) using a vibroslicer (Leica VT1200S, Leica Microsystems, Nussloch, Germany). Slices recovered for 1 h at room temperature in an artificial cerebrospinal fluid (ACSF, in mM: 124 NaCl, 5 KCl, 1.25 NaH_2_PO_4_, 26 NaHCO_3_, 10 dextrose, 1.5 MgCl_2_, and 2.5 CaCl_2_ bubbled with a mixture of 5% CO_2_ and 95% O_2_) and then transferred to a submerged recording chamber superfused with ACSF (30 ºC, 2 ml/min). Synaptic responses were evoked with 0.2 ms pulses delivered through theta glass micropipettes (TG200-4, Warner Instruments) filled with ACSF, and extracellularly recorded in the stratum radiatum of CA1. Baseline responses were recorded at 0.033 Hz using a stimulation intensity that evoked a half-maximal response, defined as the maximal response without a population spike (pop-spike). Slices were discarded if the postpike appeared in the initial rising phase, paired-pulse facilitation at a 50 ms interval was less than 10%, or the baseline was not stable. LTP was induced using theta burst stimulation (TBS; 10 trains of four pulses each at 100 Hz; 5 Hz inter-burst interval) delivered at 0.1 Hz. LTP magnitude was calculated as the average (normalized to baseline) of the responses recorded 50–60 min after conditioning stimulation.

2. *Multielectrode array (MEA) recording*

The same brain hippocampal slices were used to record both multielectrode arrays (MEA, 252 electrodes, Multichannel Systems) and local field potentials. Hippocampal slices were mounted on a MEA matrix bathed in an ACSF medium (in mM: NaCl 125, KCl 2.5, glucose 25, NaHCO_3_ 25, NaH_2_PO4 1.25, CaCl_2_ 2, and MgCl_2_ 1) at 32 °C and constantly bubbled with 95% O_2_ and 5% CO_2_. Spike sorting Spyking-circus was used, as described at (http://www.yger.net/software/spyking-circus), to detect individual neuronal action potentials (Spikes) using the default parameters that consist of four main steps: Filtering raw extracellular traces, whitening them (to remove correlated noise), clustering action potential waveforms, and fitting them on the whitened traces.

3. *Firing and burst rate recordings*

Firing rate was computed as the number of spikes divided by the recording length. Burst rate was computed analogously as the number of bursts detected by Neuroexplorer software, which is comprised of 5 parameters defining a burst: i) Maximum beginning ISI (lower inter-spike interval bound for a burst); ii) maximum end ISI (upper inter-spike interval bound for a burst); iii) minimum inter-burst interval; iv) minimum burst duration and minimum number of spikes in the burst. We defined these parameters as 0.040 s, 0.090 s, 0.120 s, 0.015 s and 3 spikes according to our visual inspection of the data. Spontaneous Activity (SA) was recorded for 10 min, following a 30-min application of picrotoxin (PTX, 100 µM).

4. *Compound muscle action potential (CMAP) recordings*

CMAPs were measured in isoflurane-anesthetized mice using an electromyographic apparatus (Keypoint, Dantec, Les Ulis, Francer AD Instruments, Oxford, UK). Briefly, the sciatic nerve was stimulated by two electrodes placed over the lumbar vertebral column. CMAP was recorded by two electrodes placed in the belly and in the tendon of the right gastrocnemius, tibialis anterior, and triceps muscles. A reference earth electrode was placed in the left gastrocnemius and connected to the electromyography apparatus (*9*).

**Biochemical and molecular analysis**

Cells and tissue homogenates were prepared in TEN buffer (10 mM Tris-HCl pH 8.0, 1 mM EDTA, 100 mM NaCl, 1% NP-40 and protein inhibitor cocktail (Roche) and sonicated for 15 s at 30% amplitude (QSonica). Protein concentration was determined by the Pierce BCA Protein Assay kit. Samples were mixed with dithiothreitol (DTT, final concentration of 0.1M) and Laemmli sample buffer and loaded onto 10% SDS-polyacrylamide gels, transferred to nitrocellulose membrane and analyzed by western blot (WB). The following antibodies and dilutions were used for WB: rabbit anti-PSD-95 (ab18258, 1:3000; Millipore), rabbit anti-Synaptophysin (1:1000; Cell signaling 4329S), mouse anti-SNAP25 (ab41455, 1:1000), mouse anti-β-actin (1:20,000, Cell Signaling 5125S), mouse anti-α-tubulin (1:5000, Millipore CP06-100), goat anti-rabbit HRP conjugate (1:2000, Invitrogen 65-6120), and goat anti-mouse HRP conjugate (1:2000, Invitrogen 62-6520). Total tissue RNA was extracted using Trizol™ Reagent (Invitrogen) according to the manufacturer’s instruction. cDNA was synthesized with random primers using a High Capacity cDNA Reverse Transcription KIT (Applied Biosystems) and subsequently subjected to quantitative PCR analysis using HOT FIREPol® EvaGreen® qPCR Mix plus (ROX) (Solis Bio Dyne) on a Stratagene Mx3000P machine (Agilent Technologies). Actin mRNA expression was used to normalize all samples. The sequences of primers used were as follows:

**Gene	Forward primer	Reverse primer**

Xbp1s	TGC TGA GTC GGC AGC AGG TG	GAC TAG CAG ACT CTG GGG AAG

Hspa5	TCA TCG GAC GCA CTT GGA A	CAA CCA CCT TGA ATG GCA AGA

Ddit3	TGG AGA GCG AGG GCT TTG	GTC CCT AGC TTG GCT GAC AGA

Ern1	CCG AGC CAT GAG AAA CAA GAA	GGG AAG CGG GAA GTG AAG TAG

Actin	TAC CAC CAT GTA CCC AGG CA	CTC AGG AGG AGC AAT GAT CTT

Atf3	TTCTTGTTTCGACACTTGGCA	CAGACCCCTGGAGATGTCAGT

**Detection of S-nitrosylated (SNO-)IRE1α**

We performed biotin-switch assays using whole-brain tissue samples as previously described with minor modifications *(10).* Briefly, brain tissue extracts were prepared in 400 μl HEN-RIPA buffer (100 mM Hepes pH 7.5, 1 mM EDTA, 0.1 mM neocuproine, 150 mM NaCl, 1% NP-40, 0.5% sodium deoxycholate, and 0.1% SDS) containing a thiol blocking reagent (20 mM methyl methanethiosulfonate [MMTS]) in a 3 ml homogenizer. After centrifuging to remove tissue debris, SDS was added to each sample to a final concentration of 1%. Samples were then incubated for 30 min at 42 °C to block free thiol groups. After removing excess MMTS by acetone precipitation, S-nitrosothiols were reduced to thiols with 10 mM ascorbate. The newly formed thiols were then linked with the sulfhydryl-specific biotinylating reagent N-[6-(biotinamido)hexyl]-3’-(2’-pyridyldithio)propionamide (Biotin-HPDP). The biotinylated proteins were pulled down with Streptavidin-agarose beads, and western blot analysis performed to detect SNO-IRE1α (1:1000, Cell Signaling 3294S). To monitor the amount of “input” protein by immunoblot, a 20 µl aliquot of the sample was saved prior to the step of Streptavidin-agarose bead addition in the biotin-switch assay. GAPDH was monitored as a control to ensure equal loading (1:1000, Millipore MAB374).

**Quantitative proteomic analysis**

The hippocampi of mice transduced with AAV2 (Mock or XBP1s) were homogenized in TEN buffer (10 mM Tris-HCl pH 8.0, 1 mM EDTA, 100 mM NaCl, 1% NP-40) and protein inhibitor cocktail (Roche), and then sonicated for 15 s at 30% amplitude (QSonica). For each sample lysate, 20 μg were precipitated with chloroform/methanol. Samples for mass spectrometry analysis were prepared as described (*11*). Air-dried pellets were resuspended in 1% RapiGest SF (Waters) and diluted to final volume in 100 mM HEPES (pH 8.0). Proteins were reduced with 5 mM Tris(2-carboxyethyl)phosphine hydrochloride (Thermo Fisher) for 30 min and alkylated with 10 mM iodoacetamide (Sigma Aldrich) for 30 min at room temperature in the dark. Proteins were then digested for 18 h at 37 °C with 0.5 μg trypsin (Promega). After digestion, the peptides from each sample were reacted for 1 h with the appropriate tandem mass tag (TMT) isobaric reagent (Thermo Fisher) in 40% (v/v) anhydrous acetonitrile and quenched with 0.4% ammonium bicarbonate for 1 h. Samples with different TMT labels were pooled and acidified with 5% formic acid. Acetonitrile was evaporated on a SpeedVac and debris removed by centrifugation for 30 min at 18,000g. MudPIT microcolumns were prepared as described (*12*). Liquid chromatography-tandem mass spectrometry (LC-MS/MS) analysis was performed using a Q-Exactive HF mass spectrometer equipped with an Ultimate 3000 nLC 1000 (Thermo Fisher). MudPIT experiments were performed by 10 μl sequential injections of 0, 10, 20, 30, . . . , 100% buffer C (500 mM ammonium acetate in buffer A) and a final step of 90% buffer C/10% buffer B (100% acetonitrile, 0.1% formic acid, v/v/v), with each step followed by a gradient from buffer A (95% water, 5% acetonitrile, 0.1% formic acid) to buffer B. Electrospray was performed directly from the analytical column by applying a voltage of 2 kV with an inlet capillary temperature of 275 °C. Data-dependent acquisition of MS/MS spectra was performed with the following settings: Eluted peptides were scanned from 300 to 1800 m/z with a resolution of 120,000. The top 15 peaks for each full scan were fragmented by higher energy collisional dissociation (HCD) using a normalized collision energy of 38%, isolation window of 0.7 m/z, a resolution of 45,000, ACG target 1e5, maximum IT 60 ms, and scanned from 100 to 1800 m/z. Dynamic exclusion was set to 10 s. Peptide identification and protein quantification was performed using Proteome Discoverer 2.2 (ThermoFisher). Spectra were searched using SEQUEST against a UniProt mouse proteome database. The database was curated to remove redundant protein and splice-isoforms, and common contaminants were added. Searches were carried out using a decoy database of reversed peptide sequences using Percolator node for filtering and the following settings: 10 ppm peptide precursor tolerance, 6 amino acid minimum peptide length, trypsin cleavage (maximum 2 missed cleavage events), static Cys modification of 57.021517 (carbamidomethylation), and static N-terminal and Lys modification of 229.1629 (TMT-sixplex), FDR 0.01, 2 peptide IDs per protein. Normalization of TMT reporter ion intensities was carried out based on total peptide abundance in each channel, and subsequently, TMT ratios for each identified protein were calculated in reference to a common pooled sample. Finally, the reference-normalized TMT intensities were compared between young WT vs.XBP1s Tg (*n* = 4, 4), middle-aged WT vs. XBP1s Tg (*n* = 3, 4), and aged WT vs. XBP1s Tg (*n* = 3, 4), with transduced samples and significance assessed by a two-tailed unpaired Student’s t-test (*13*) and Q = 1% in Graphpad Prism. Enrichment analysis based on the functional annotation of the differentially expressed genes (*P* < 0.05) was performed using the Kyoto Encyclopedia for Genes and Genomes (KEGG), Gene Ontology, and TRRUST v2 terms through the EnrichR platform or STRING v.11. Protein-protein association networks were generated with STRING v.11 (*14, 15*). Heatmaps were generated using Morpheus (<https://software.broadinstitute.org/morpheus>).

**Neuromuscular junction (NMJ) staining and analyses**

The levator auris longus (LAL) muscles were dissected and whole-mount fixed in 0.5% formaldehyde (FA) for 90 min. Samples were blocked with 4% bovine serum albumin (BSA) dissolved in PBST 12-16 h at 4 °C. Muscles were incubated with mouse monoclonal antibodies against neurofilament (2H3, 1:300) and synaptic vesicles (SV2, 1:50; both from the Developmental Studies Hybridoma Bank, DSHB, of the University of Iowa, USA) in 4% BSA-PBST for 30 min at RT and then 12-16 h at 4 °C. Tissues were then incubated with the respective secondary antibodies (1:300, Jackson Immuno Research, West Grove, PA, USA) in 4% BSA-PBST containing Alexa488-conjugated α-bungarotoxin (BTX, 1:500, Invitrogen, Carlsbad, CA, USA) and DAPI (1:1000, Thermo Fisher, Waltham, MA, USA) for 12-16 h at 4 °C. Samples were post-fixed with 1% FA in 1X PBS for 10 min at 22 °C, flat mounted and imaged. Z-stack images were collected at 1-μm intervals on a Zeiss LSM 700 confocal microscope. Maximal intensity projection images were reconstructed in 3D using ImageJ software. The morphology of >50 NMJs per mouse was manually determined and expressed as a percentage of the total. The area of >50 acetylcholine receptor (AChR) densities per mouse was determined for each postsynaptic structure using ImageJ software.

**Statistical analysis**

Statistical significance respective to age was queried for single comparisons with a Student’s t test and for multiple comparisons with a one-way ANOVA followed by Tukey’s post-hoc test. When analyzing both age and genotype in the same comparison, a two-way ANOVA followed by Tukey’s post-hoc test was implemented. *P* values were considered significant when they were < 0.05. The number of animals in each group varied from 4-15 depending on the experiment based on a Power Analysis of prior data.

**Materials and Methods References**

1. P. Valdés, G. Mercado, R.L. Vidal, C. Molina, G. Parsons, A. Martinez,... C. Hetz, Control of dopaminergic neuron survival by the unfolded protein response transcription factor XBP1. *Proc. Natl. Acad. Sci. U.S.A* **111.18** 6804-6809 (2014).
2. M. Leger, A. Quiedeville, V. Bouet, B. Haelewyn, M. Boulouard, P. Schumann-Bard, T. Freret, Object recognition test in mice. *Nat. Protoc* **8.12** 2531 (2013).
3. A. Vogel-Ciernia, M.A. Wood, Examining object location and object recognition memory in mice. *Curr Protoc Neurosci* **69.1** 8-31 (2014).
4. C.A. Barnes, Memory deficits associated with senescence: a neurophysiological and behavioral study in the rat. *J Comp Physiol Psychol*. **93** 74-104 (1979).
5. B. Sunyer, S. Patil, H. Höger, G. Luber, Barnes maze, a useful task to assess spatial reference memory in the mice. *Nat Protoc* **390** 10-38 (2007).
6. F. Debacq-Chainiaux, J.D. Erusalimsky, J. Campisi, O. Toussaint, Protocols to detect senescence-associated beta-galactosidase (SA-betagal) activity, a biomarker of senescent cells in culture and in vivo. *Nat Protoc* **4** (12) 1798-1806 (2009).
7. A.O. Ardiles, C.C. Tapia-Rojas, M. Mandal, F. Alexandre, A. Kirkwood, N.C. Inestrosa, A.G. Palacios, Postsynaptic dysfunction is associated with spatial and object recognition memory loss in a natural model of Alzheimer’s disease. *Proc. Natl. Acad. Sci. U.S.A* , ***109*(34)** 13835-13840 (2012).
8. C.R. Ravello, L.U. Perrinet, M.J. Escobar, A.G. Palacios, Speed-selectivity in retinal ganglion cells is sharpened by broad spatial frequency, naturalistic stimuli. *Sci. Rep.* ***9*(1)**, 1-16 (2019).
9. E. Dirren, J. Aebischer, C. Rochat, C. Towne, B.L. Schneider, P. Aebischer, SOD1 silencing in motoneurons or glia rescues neuromuscular function in ALS mice. *Ann. Clin. Transl. Neurol.*, ***2*(2)** 167-184 (2015).
10. T. Uehara, T. Nakamura, D. Yao, Z.Q. Shi, Z. Gu, Y. Ma, , ... S.A. Lipton, S-nitrosylated protein-disulphide isomerase links protein misfolding to neurodegeneration. *Nature*, ***441*(7092),** 513-517 (2006).
11. L. Plate, C.B. Cooley, J.J. Chen, R.J. Paxman, C.M. Gallagher, F. Madoux, L. Scampavia, Small molecule proteostasis regulators that reprogram the ER to reduce extracellular protein aggregation. *Elife*, ***5*,** e15550 (2016).
12. L.M. Ryno, J.C. Genereux, J. C., Naito, T., Morimoto, R. I., Powers, E. T., Shoulders, M. D., Wiseman, R. L. Characterizing the altered cellular proteome induced by the stress-independent activation of heat shock factor 1. *ACS Chem. Biol*, ***9*(6**), 1273-1283. (2014).

13. Y. Benjamini, A.M. Krieger, D. Yekutieli, Adaptive linear step-up procedures that control the false discovery rate. *Biometrika* **93**, 491–507 (2006).

14. M.V. Kuleshov, M.R. Jones, A.D. Rouillard, N.F. Fernandez, Q. Duan, Z. Wang,... M.G. McDermott, Enrichr: a comprehensive gene set enrichment analysis web server 2016 update. *Nuc Acids Res*, **44(W1)**, W90-W97 (2016).

15.	D. Szklarczyk , A.L. Gable, D. Lyon, A. Junge, S. Wyder , J. Huerta-Cepas, M. Simonovic, N.T. Doncheva, J.H. Morris, P. Bork,L.J. Jensen, STRING v11: protein–protein association networks with increased coverage, supporting functional discovery in genome-wide experimental datasets. *Nuc Acids Res* **47(D1)**: D607-13 (2018).

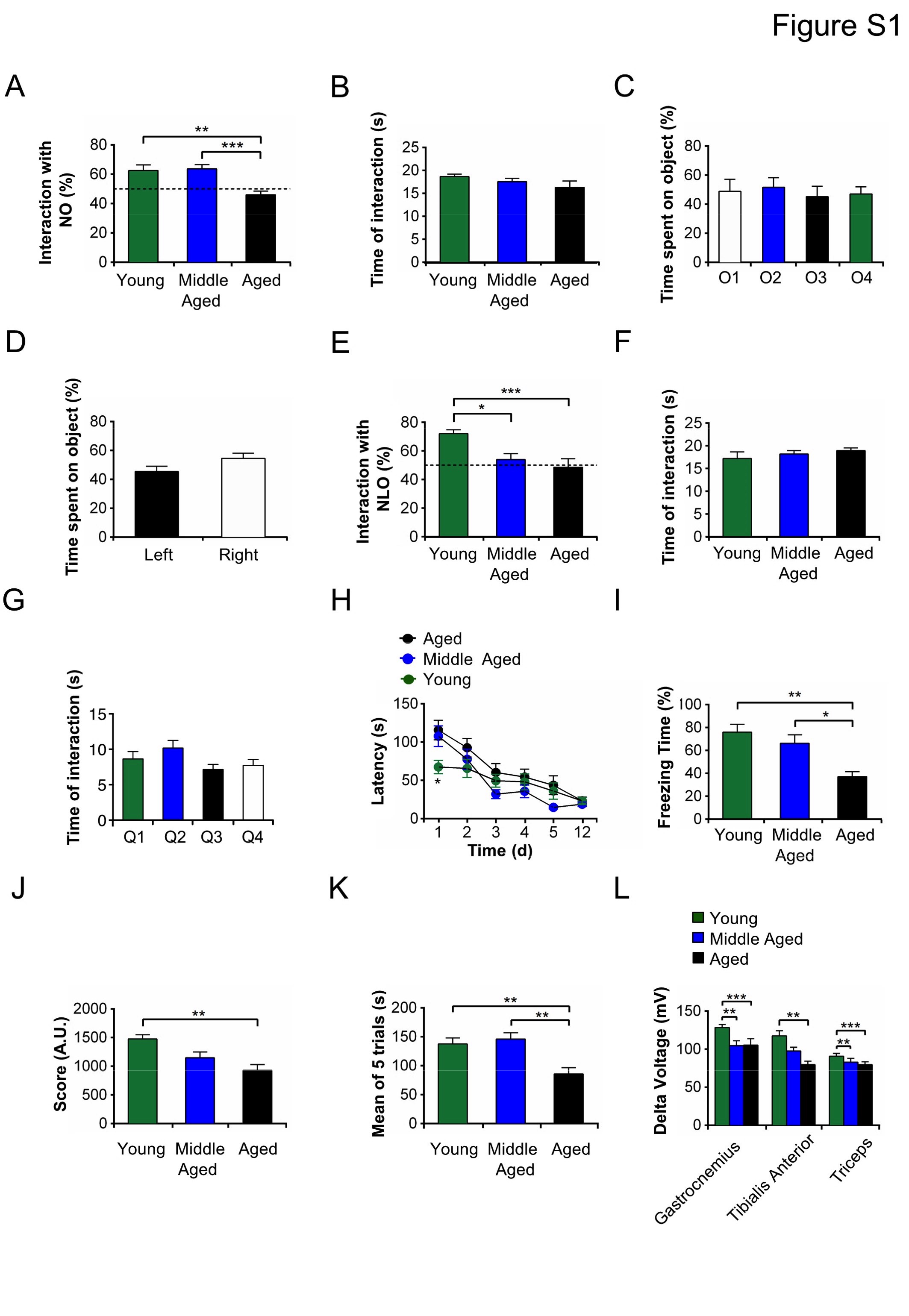

**Fig. S1.**

**Figure Supplementary 1.** Functional tests for age-associated cognitive and motor decline. (**A**) Young, middle-aged, and aged WT animals were evaluated on the new object recognition test, comparing the new/total interaction time with novel objects following 24 h of presentation of two identical objects; this tested the ability of the mice to discriminate novel objects. Histogram shows mean and SEM of percentages of interaction time with novel objects (*n* = 14, 17, 18 for the various ages, respectively). ***P* < 0.01, ****P* < 0.005 by one-way ANOVA followed by Tukey’s post-hoc test. (**B**) Young, middle-aged, and aged WT animals were evaluated on the first day of the new object recognition (NOR) test to compare the total time of interaction with any of the objects presented. Maximum time of interaction was set at 5 minutes. If animals reached the criteria to interact for at least 20 seconds with both objects, the test was completed. Histogram show mean and SEM of interaction time with any object (*n* = 40, 29, 21 for the various ages, respectively). (**C**) Young, middle-aged, and aged WT animals were introduced to the four different types of objects implemented in the NOR test. Histogram shows mean and SEM of total time of interaction with each type of object (*n* = 9, 8, 9, 10 for each object). (**D**) Young, middle-aged, and aged animals were allowed to interact with identical objects in the two different quadrants of the arena used in the NOR test (left or right). Histogram shows mean and SEM of time spent on objects placed in different quadrants in the cage (n = 14, 14/quadrant). (**E**) Young, middle-aged, and aged WT animals were evaluated in the new object location test comparing the percentage of interaction time with novelly-located object (NLO) following 24 h of presentation to one object placed in a different position. Histogram shows mean and SEM of percentages of interaction time with the novelly-located object (*n* = 18, 13, 17 for the various ages, respectively). **P* < 0.05, ****P* < 0.005 by one-way ANOVA followed by Tukey’s post-hoc test. (**F**) Young, middle-aged, and aged WT animals were evaluated on the first day of the NOL test to compare the total time of interaction with any of the objects presented. Maximum time of interaction was set at 5 minutes. If an animal reached the criteria to interact of at least 20 seconds with both objects, the test was considered completed. Histogram shows mean and SEM of the sum of interaction time with any object (*n* = 19, 20, 19 for the various ages, respectively). (**G**) Young, middle-aged, and aged WT animals were allowed to interact with identical objects in the 4 different quadrants of the arena used in the NOL test (quadrant 1, 2, 3 or 4). Histogram shows mean and SEM of time spent with objects placed in different quadrants in the cage used in the test (*n* = 17-23 animals/group). (**H**) Young, middle-aged, and aged WT animals were evaluated in the Barnes maze test to measure spatial memory ability. Graph shows mean and SEM of primary latency to reach the target hole on each day of the test (*n* = 10/group). **P* < 0.5 by Student’s t test with Bonferroni correction performed on each day. **(I)** Young, middle-aged, and aged WT animals were evaluated in the contextual fear conditioning test 24 h after presentation of an unconditioned stimulus. Histogram shows mean and SEM of percentage of freezing time (*n* = 6/group). **P* < 0.05, ***P* < 0.001 by one-way ANOVA followed by Tukey’s post-hoc test). (**J**) Young, middle-aged, and aged WT animals were evaluated in the wire hanging test with a score based on total time of hanging and falls off of the wire in a 3 min period. Histogram shows mean and SEM of score in arbitrary units (au; *n* = 9, 11, 15 for each age group, respectively). ***P* < 0.01 by one-way ANOVA followed by Tukey’s post-hoc test. (**K**) Young, middle-aged, and aged WT animals were evaluated in the rotarod test to analyze muscular function and coordination. Histogram shows mean and SEM of latencies to fall from the rod under constant acceleration in 4 trials split over 4 consecutive days (*n* = 16, 8, 17 for each age group, respectively). ***P* < 0.01 by one-way ANOVA followed by Tukey’s post-hoc test. **(L)** Young, middle-aged, and aged WT animals were deeply anesthetized and had their compound muscle amplitude potentials (CMAPs) measured in the gastrocnemius, tibialis anterior, and triceps muscles. Histogram shows mean and SEM of differences between maximum and minimum amplitudes (*n* = 8, 8, 8 for each age group, respectively). ***P* < 0.01, ****P* < 0.001 by one-way ANOVA followed by Tukey’s post-hoc test.

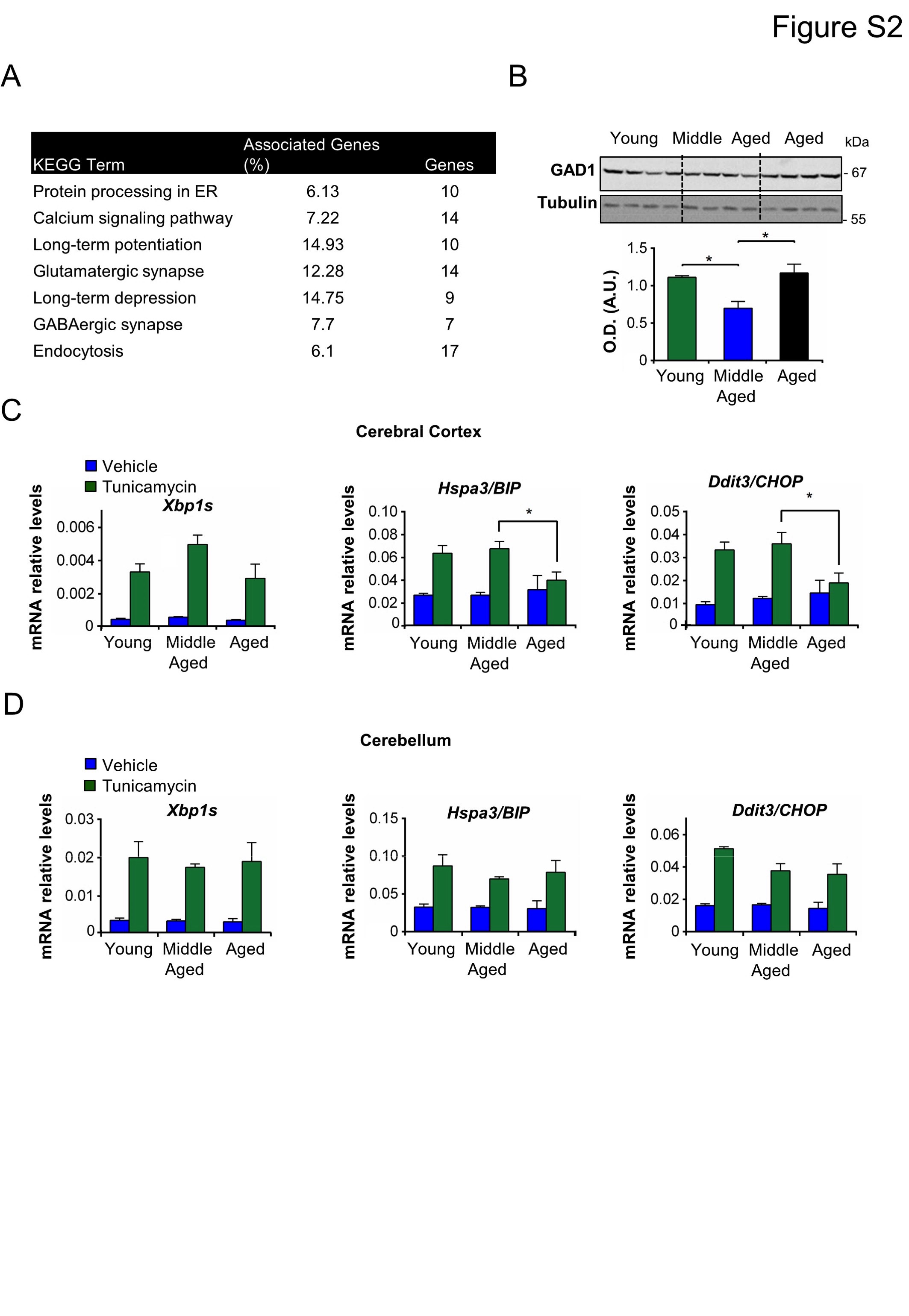

**Figure Supplementary 2. (A)** Most significant proteomic changes comparing hippocampi of young vs. middle-aged animals with cut-off of *P* < 0.05 (*n* = 4 animals/group). Functional enrichment analysis on KEGG database using EnrichR platform was performed with major enriched terms depicted. **(B)** Western blots comparing total hippocampal extracts of young, middle-aged, and aged WT animals for protein levels of GAD-1, a major XBP-1s target gene found in the proteomics analysis. Histogram shows mean and SEM for densitometry of each protein band in arbitrary units (A.U.) normalized to tubulin protein levels as a loading control (*n* = 4/group). **P* < 0.05 by one-way ANOVA followed by Tukey’s post-hoc test. **(C** and **D)** Young, middle-aged, and aged WT animals were treated with tunicamycin (5 mg/kg) or vehicle for 24 h. Following euthanasia, the cerebrocortex and cerebellum were dissected for RNA extraction. Histograms show mean and SEM of qPCR for *Xbp1s, Hspa5/Bip*, and *Ddit3/Chop* relative mRNA levels in hippocampal samples from treated animals compared to controls (ˆ = 4 animals/group). **P*< 0.05 by one-way ANOVA followed by Tukey’s post-hoc test on treated groups.

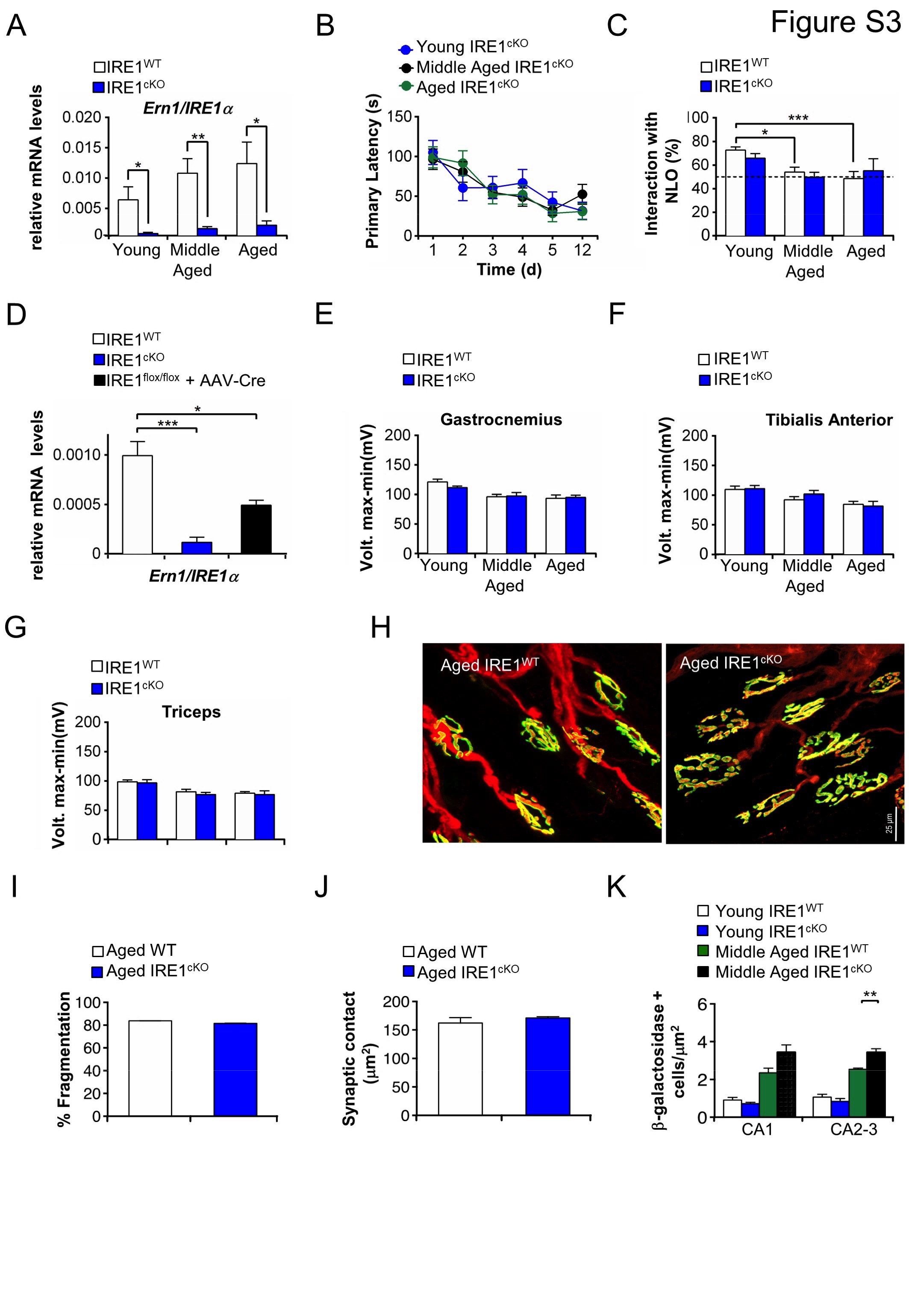

**Figure Supplementary 3.** (**A**) Relative mRNA levels of *Ern1* sequence (exon 20-21) by qPCR from hippocampi of young, middle-aged, and aged IRE1^flox/flox^ or IRE1^cKO^ animals (*n* = 4 per genotype at each age). ***P* < 0.01 by unpaired Student’s t test for each age group). (**B**) Young, middle-aged, and aged IRE1^cKO^ were trained over a period of 5 days on the Barnes maze, a test of spatial learning and memory. One week later, animals were re-evaluated (on day 12) to measure consolidation of long-term spatial memory. Graphs shows mean and SEM of primary latency to reach the target hole on a given day (n = 10, 10, 8 for each age group, respectively). (**C**) New object location test to evaluate the ability to discriminate changes in object location 24 h after presentation of two displaced objects to IRE1^WT^ and IRE1^cKO^ animals at young (*n* = 17, 21, respectively, by genotype), middle-aged (*n* = 13, 13), or aged (*n* = 17, 8) time points. Histogram shows mean and SEM of interaction time with the novelly-located object (NLO). *P* < 0.05, ****P* < 0.005 comparing age-matched groups by one-way ANOVA followed by Tukey’s post-hoc test. (**D**) qPCR of relative mRNA levels of *Ern1* sequence exon 20-21 in hippocampi of middle-aged IRE1^flox/flox^, IRE1^cKO^, and IRE1^flox/flox^ mice injected with AAV2-CRE (*n* = 4 animals/group). **P* < 0.05, ****P* < 0.005 by one-way ANOVA followed by Tukey’s post-hoc test. (**E-G**) Compound muscle amplitudes potentials (CMAPs) of various muscles from young, middle-aged, and aged IRE1^WT^ or IRE1^cKO^ mice. Histograms show mean and SEM for difference between the minimum and maximum amplitudes (*n* = 10-11 animals/group). There was no statistical difference by unpaired Student’s t test within each age-matched group. (**H-J**) Morphological analysis of neuromuscular junctions (NMJs) of levator auris longus muscle in elderly mice from aged IRE1^WT^ or IRE1^cKO^ animals. After immunolabeling, NMJs were imaged by confocal microscopy (40x magnification; scale bar, 25 m). Representative images for aged WT and aged IRE1^cKO^ are shown **(H)**. Histograms show mean and SEM of percentage of fragmentation **(I)** and synaptic contact of NMJs **(J)** (*n* = 4 animals/group). **(K)** -Galactosidase positive cells were measured in the hippocampus of young and middle-aged IRE1^WT^ or IRE1^cKO^ mice. Histogram shows mean and SEM of positively-labeled cells in the hippocampus (*n* = 3-4 animals/group). ***P* < 0.01 by unpaired Student’s t test

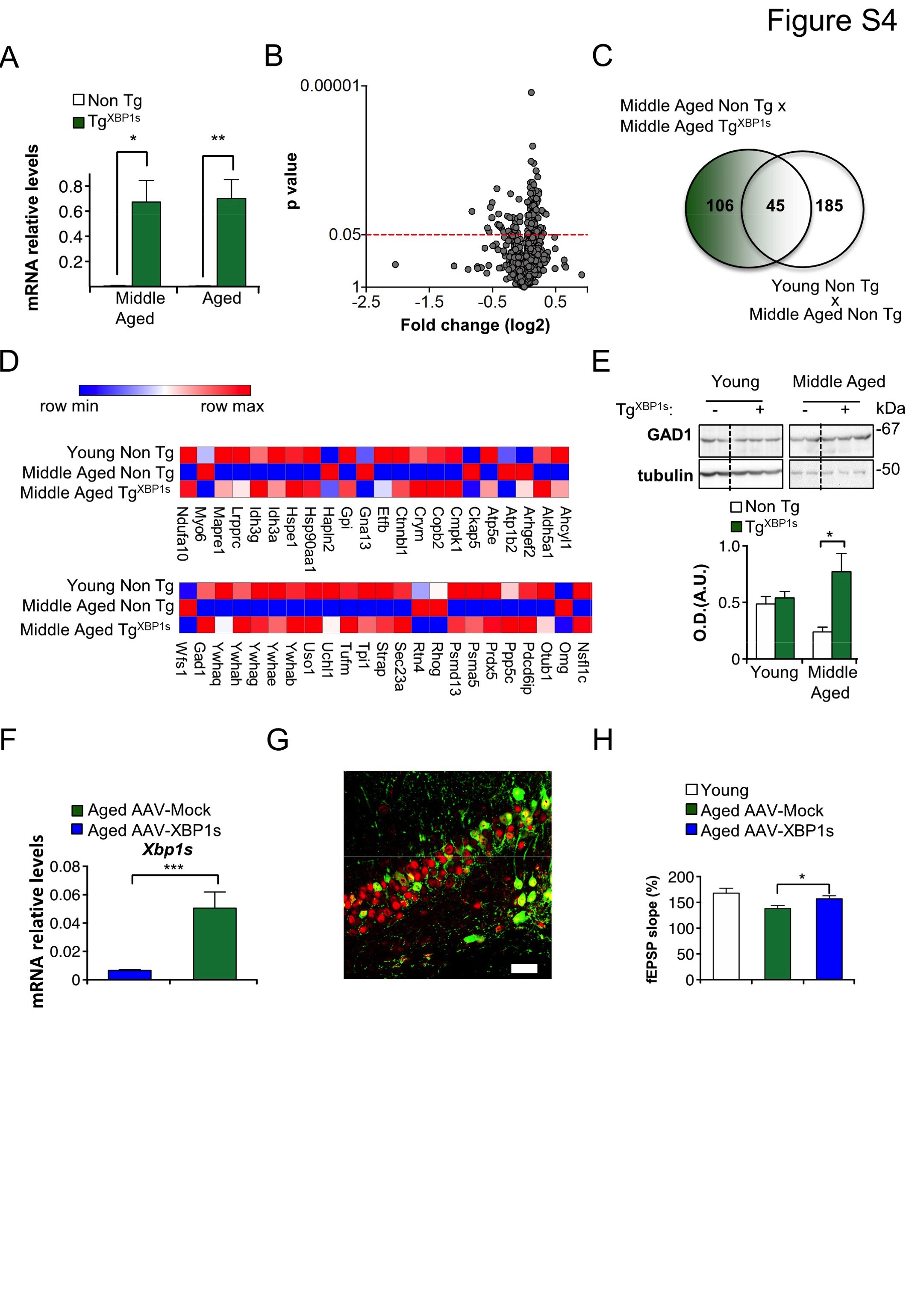

**Figure Supplementary 4.** (**A**) Relative mRNA levels of *Xbp1s* by qPCR in middle-aged and aged non-Tg or Tg^XBP1s^ hippocampi (*n*=3-6 animals/group). **P* < 0.05, ***P* < 0.01 within age-matched groups by unpaired Student’s t test). (**B**) Volcano plot for proteomic comparison of hippocampal tissue derived from middle-aged non-Tg and middle-aged Tg^XBP1s^ animals indicating *P* values (y-axis) and fold-change (log2) (x-axis). Red dotted lines delineate cut-off used to filter genes for functional enrichment analysis. (**C**) Venn diagram showing number of significant proteomic alterations (cut-off: *P* < 0.05) comparing young non-Tg, middle-aged non-Tg ,and middle-aged Tg^XBP1s^ mice (*n* = 3-4 animals/group). Stop **(D)** Heat maps indicating significant gene alterations among young non-Tg, middle-aged non-Tg, and middle-aged Tg^XBP1s^ mice. **(E)** Western blots of GAD-1 comparing total hippocampal extracts of young and middle-aged non-Tg or Tg^XBP1s^. Upper panel: Representative gels. Lower panel: Histogram showing mean and SEM optical density quantified in arbitrary units (A.U) for each sample normalized to tubulin as loading control (*n* = 4/group). **P* < 0.05 by unpaired Student’s t test. **(F)** qPCR of relative mRNA levels of *Xbp1s* in hippocampi of aged mice injected with AAV2-XBP1s or AAV2-Mock control. **G**) Representative confocal photomicrograph of immunolabel for NeuN (red) and GFP (green) from CA1 region of hippocampus of aged mouse brain 4 weeks after injection with AAV-XBP1s. (**H**) Field excitatory postsynaptic potential (fEPSP) slopes obtained 1 h after induction of LTP in hippocampal slices of aged mice previously injected with AAV2-Mock or AAV2-XBP1s (*n* = 6, 9 animals with 17-28 slices/animal). **P* < 0.05 by unpaired Student’s t test. fEPSP from young animals shown for comparison.

**Table S1.**

Hippocampal proteome derived from young and middle-aged WT animals. Statistically significant alterations (*P* < 0.05) are highlighted. Table shows genes encoding the proteins, differences in expression levels, and *P* values (*n* = 4 animals/group).

| **Gene** | **Difference** | **p Value** |
| --- | --- | --- |
| Lrpprc | -0.190936724 | 1.06982E-06 |
| Dpysl5 | -0.460324157 | 2.48756E-05 |
| Cacng8 | 0.468115181 | 4.71233E-05 |
| Shank1 | 0.24176101 | 7.05805E-05 |
| Arhgef2 | 0.180347143 | 8.23916E-05 |
| Calb2 | -1.693371991 | 8.44896E-05 |
| Shisa7 | 0.37529194 | 0.00016748 |
| Dlg4 | 0.208689771 | 0.000212352 |
| Abat | -0.23280303 | 0.000235396 |
| Kif1a | -0.739707291 | 0.000287857 |
| Psma5 | -0.305528878 | 0.000333731 |
| Anks1b | 0.294753529 | 0.000367142 |
| Pclo | 0.201361633 | 0.000443373 |
| Shisa6 | 0.324770913 | 0.000470047 |
| Syngap1 | 0.314772466 | 0.000591216 |
| Scrn3 | -0.099392347 | 0.000593922 |
| Grin2a | 0.325955858 | 0.000662481 |
| Gria1 | 0.336342117 | 0.001067323 |
| Cltb | -0.436801579 | 0.001074668 |
| Gmps | -0.203980528 | 0.001107336 |
| Cct8 | -0.126755968 | 0.001158932 |
| Camk2a | 0.265462678 | 0.001178181 |
| Copb2 | -0.206023903 | 0.001320536 |
| Dpp6 | 0.202983637 | 0.001371231 |
| Map6d1 | 0.358647132 | 0.001414177 |
| Capza2 | 0.140530703 | 0.001570731 |
| Dctn1 | -0.063269601 | 0.001589871 |
| Slc6a11 | -0.421828591 | 0.001645415 |
| Hapln2 | 1.068256898 | 0.001668986 |
| Iqsec2 | 0.259245371 | 0.00177361 |
| Baiap2 | 0.154222358 | 0.002238798 |
| Psd | 0.333064786 | 0.002294109 |
| Gda | -0.335973187 | 0.002390571 |
| Psmd5 | -0.199424872 | 0.002446258 |
| Slc44a2 | 0.341211531 | 0.002557812 |
| Prkcg | 0.20972723 | 0.00269638 |
| Atp2b2 | 0.134473622 | 0.002773552 |
| Rhog | 0.262153062 | 0.003290931 |
| Cntnap1 | 0.2481266 | 0.003420951 |
| Ccar2 | -0.216750395 | 0.003458747 |
| Alb | -0.69106096 | 0.003535914 |
| Ahcyl1 | -0.114447079 | 0.003818183 |
| Iqsec1 | 0.143992143 | 0.003882748 |
| Gria2 | 0.222842894 | 0.004043964 |
| Idh3g | -0.103943103 | 0.004085888 |
| Icam5 | 0.260571714 | 0.004203034 |
| Prpsap2 | 0.414972283 | 0.004323684 |
| Arpc5 | 0.322281573 | 0.004368072 |
| Set | -0.250854262 | 0.004480596 |
| Homer1 | 0.133998745 | 0.004507935 |
| Psmd13 | -0.126072034 | 0.004533275 |
| Cttn | 0.167346838 | 0.004581382 |
| Snx1 | -0.284990233 | 0.004771495 |
| Glud1 | -0.095729105 | 0.005141298 |
| Dnm3 | -0.183930799 | 0.005282096 |
| Phb | -0.113905153 | 0.005286205 |
| Ywhab | -0.16834362 | 0.005306849 |
| Dpysl3 | -0.27666012 | 0.00534771 |
| Stk39 | -0.29874131 | 0.005372366 |
| Dlgap3 | 0.181292293 | 0.00562536 |
| Camk2b | 0.198014531 | 0.005767375 |
| Prdx2 | -0.163749495 | 0.005792336 |
| Otub1 | -0.363697416 | 0.00600267 |
| Kalrn | 0.420708346 | 0.006104069 |
| Aldh2 | -0.157532833 | 0.006107051 |
| Idh3a | -0.090607765 | 0.006216212 |
| Erc2 | 0.205861752 | 0.006297527 |
| Grin1 | 0.204059013 | 0.006476326 |
| Nfasc | 0.181035435 | 0.006624677 |
| Mgea5 | -0.193861288 | 0.006723781 |
| Stmn1 | -0.528096141 | 0.006768116 |
| Arf3 | 0.207834918 | 0.006856808 |
| Dync1h1 | -0.174188056 | 0.007157781 |
| Osbpl1a | -0.307945135 | 0.007336956 |
| Tuba4a | 0.218975145 | 0.007517857 |
| Ywhae | -0.299629839 | 0.007547402 |
| Rtn4 | 0.070179382 | 0.007569793 |
| Cox5a | -0.744184049 | 0.007682472 |
| Atp5e | -0.079774816 | 0.007720774 |
| Prkar2b | -0.248022031 | 0.007941743 |
| Ywhaq | -0.300236902 | 0.007964824 |
| Kcnd2 | 0.249174826 | 0.008355463 |
| Akr1a1 | -0.106258468 | 0.008363987 |
| Ndufb10 | 0.236223201 | 0.008395011 |
| Uchl1 | -0.390745983 | 0.009038767 |
| Itpr1 | 0.152126925 | 0.009080007 |
| Cend1 | -0.127581802 | 0.009325072 |
| Tpi1 | -0.119557114 | 0.009747851 |
| Gap43 | -0.418597961 | 0.009995776 |
| Lin7b | 0.190642979 | 0.009996325 |
| Ttyh1 | 0.256387039 | 0.010148568 |
| Crym | -0.200181651 | 0.010675534 |
| Pak1 | -0.193513814 | 0.010867588 |
| Cct5 | -0.106312217 | 0.01117054 |
| Dnajb6 | 0.15092962 | 0.011504632 |
| Uso1 | -0.179899872 | 0.01159842 |
| Gad1 | -0.129022631 | 0.011898917 |
| Dlg3 | 0.151356399 | 0.012008946 |
| Sec23a | -0.126878449 | 0.012300419 |
| Epha4 | 0.262099745 | 0.012938893 |
| Arpc2 | 0.197196028 | 0.013059562 |
| Ywhag | -0.249682869 | 0.013334746 |
| Ablim2 | 0.198417986 | 0.013531689 |
| Hsp90ab1 | -0.192440591 | 0.013754967 |
| Pgm1 | -0.152991135 | 0.013984187 |
| Cmpk1 | -0.182456803 | 0.014676463 |
| Ran | -0.12184637 | 0.015104135 |
| Ahcyl2 | -0.26967135 | 0.015125748 |
| Fubp1 | -0.231132416 | 0.015182452 |
| Mlf2 | 0.170291009 | 0.015276099 |
| Acaa2 | -0.377502968 | 0.015299022 |
| Sh3glb2 | 0.102128647 | 0.015776151 |
| Eef1g | -0.18254811 | 0.015903846 |
| Hint2 | -0.221021867 | 0.016356217 |
| Ckap5 | 0.2208133 | 0.016524919 |
| Slc4a10 | -0.264014495 | 0.016559511 |
| Idh2 | -0.170202841 | 0.01659275 |
| Stxbp5l | 0.161706908 | 0.016685873 |
| Mapre1 | -0.175589346 | 0.01687275 |
| C1qa | 0.69323415 | 0.016881196 |
| Ivd | -0.222901369 | 0.016891404 |
| S100a1 | -1.330491662 | 0.017419567 |
| Uqcrh | -0.627930288 | 0.018192818 |
| Lppr4 | 0.125089334 | 0.018502316 |
| Cnksr2 | 0.230191371 | 0.018917894 |
| Shank3 | 0.195243936 | 0.01896023 |
| Hspa12a | -0.060768172 | 0.019258104 |
| Aldh7a1 | -0.198187432 | 0.019422709 |
| Agap2 | 0.128583906 | 0.019564164 |
| Khsrp | -0.161365749 | 0.019823354 |
| Hsp90aa1 | -0.182244625 | 0.020017859 |
| Etfa | -0.21075486 | 0.020946195 |
| Gnaq | 0.228164232 | 0.021046081 |
| Kbtbd11 | 0.112427142 | 0.021463093 |
| Hsp90b1 | -0.239360049 | 0.021759473 |
| Sf3b3 | -0.186801581 | 0.021833163 |
| Vdac2 | -0.192597592 | 0.022206879 |
| Prdx5 | -0.10950389 | 0.022385237 |
| Fh | -0.087906766 | 0.023290055 |
| Pdcd6ip | -0.132413226 | 0.02377646 |
| Adss | -0.143930891 | 0.024179766 |
| Calb1 | -0.559751863 | 0.024353473 |
| Cacna2d1 | 0.118235513 | 0.024531797 |
| Ppp5c | -0.10278737 | 0.024548367 |
| Snap25 | 0.140263478 | 0.024874866 |
| Adam10 | 0.339597909 | 0.02507354 |
| Gpi | -0.088329731 | 0.025218739 |
| Ywhah | -0.126380739 | 0.02545788 |
| Glod4 | -0.192928749 | 0.025633984 |
| Pde1b | -0.228547011 | 0.026345252 |
| Omg | 0.218252605 | 0.026431647 |
| Prepl | -0.200063221 | 0.026442551 |
| Pgrmc1 | -0.281512024 | 0.026444455 |
| P4hb | -0.192250153 | 0.026459989 |
| Cacna1a | 0.141560915 | 0.026813546 |
| Dlg2 | 0.184417178 | 0.026831172 |
| Cct3 | -0.108084253 | 0.027097121 |
| Ndufv2 | -0.206266129 | 0.027246524 |
| Gdi1 | -0.10846319 | 0.02731653 |
| Etfb | -0.172405428 | 0.027604726 |
| Pde1a | -0.285493001 | 0.02760618 |
| Pdia6 | -0.241182525 | 0.027692325 |
| Maoa | -0.148850498 | 0.027856358 |
| Ksr1 | 0.201731836 | 0.027887564 |
| Gna11 | 0.157742135 | 0.028279152 |
| Ranbp1 | -0.315796345 | 0.028318881 |
| Syt7 | 0.240059558 | 0.028795397 |
| Gfap | 0.749289017 | 0.02894084 |
| Cirbp | 0.278615609 | 0.029277044 |
| Syngr3 | 0.093488609 | 0.029527868 |
| Aldh18a1 | -0.176578666 | 0.029679234 |
| Srgap3 | 0.279848086 | 0.029708636 |
| Ctnnbl1 | -0.381248112 | 0.02989418 |
| Aldh5a1 | -0.125104286 | 0.030342382 |
| Rapgef2 | 0.104988991 | 0.030460221 |
| Huwe1 | -0.157716351 | 0.031070045 |
| Timm10 | -0.36393957 | 0.031089801 |
| Lasp1 | -0.19260216 | 0.031094149 |
| Hadh | -0.199229441 | 0.031346082 |
| Rab11b | -0.062082364 | 0.031367368 |
| Dlgap1 | 0.214172033 | 0.03137901 |
| Hspe1 | -0.153589572 | 0.031491132 |
| Eftud2 | 0.119228792 | 0.032015085 |
| Wfs1 | 0.29450642 | 0.033193999 |
|  | -0.255096077 | 0.033336062 |
| Cbr1 | -0.121006436 | 0.033439383 |
| Gabra1 | 0.128869872 | 0.033524574 |
| Cyb5b | 0.296076827 | 0.033632257 |
| Nap1l4 | -0.154284423 | 0.033667878 |
| Prps1 | 0.298121096 | 0.033830513 |
| Prkca | 0.168032189 | 0.033977752 |
| Hadha | -0.128064348 | 0.034392182 |
| Slc1a4 | 0.279518657 | 0.034401386 |
| Wipf3 | 0.2842958 | 0.034827289 |
| Ntrk2 | -0.337340363 | 0.034860102 |
| Nsfl1c | -0.178807808 | 0.035094853 |
| Lactb | 0.347150037 | 0.035616121 |
| Tppp3 | -0.202565028 | 0.036178122 |
| Actr3b | 0.217155554 | 0.03651082 |
| Pvalb | -1.260424423 | 0.037402449 |
| Atp1b2 | 0.203510499 | 0.037553493 |
| Add2 | 0.108562338 | 0.037565557 |
| Hepacam | 0.267641325 | 0.037591147 |
| Pcca | -0.113947503 | 0.037677893 |
| Arpc1a | 0.17703634 | 0.038108051 |
| Atl1 | 0.146622804 | 0.038394152 |
| Wars | -0.153041205 | 0.038537873 |
| Ezr | -0.224087415 | 0.038645199 |
| Add3 | 0.092931924 | 0.039274583 |
| Pc | -0.05399918 | 0.039701195 |
| Tufm | -0.105200898 | 0.039894194 |
| Pfdn2 | -0.185463164 | 0.040287988 |
| Sccpdh | -0.120828004 | 0.0405282 |
| Tars | -0.175396689 | 0.040530752 |
| Strap | -0.121942569 | 0.04105041 |
| Rab7a | 0.083088176 | 0.041055265 |
| Pgk1 | -0.1000193 | 0.041061961 |
| Arpc3 | 0.153067719 | 0.041609053 |
| Enoph1 | -0.110733086 | 0.043807735 |
| Vcp | -0.087097686 | 0.044321421 |
| Synpo | 0.161228801 | 0.045441041 |
| Pygm | -0.146765408 | 0.046082616 |
| Dock3 | 0.19486458 | 0.046397694 |
| Pdia3 | -0.121593666 | 0.046720322 |
| Ndufa10 | -0.059500594 | 0.046907991 |
| Myo6 | 0.129622713 | 0.047220661 |
| Atp8a1 | 0.125146093 | 0.048407757 |
| Lonp1 | -0.134158447 | 0.049186156 |
| Hspa9 | -0.093818814 | 0.050495726 |
| Cpne7 | -0.300148628 | 0.050556144 |
| Gna13 | 0.29955601 | 0.050852137 |
| Psmd1 | -0.131554029 | 0.050965542 |

**Table S2.**

Transcription regulatory network and enrichment analysis of most significant proteomic alterations (*P* < 0.05) comparing the hippocampi of young and middle-aged WT animals (*n* = 4 animals/group) using TRRUST v.2, KEGG 2019, and Gene Ontology 2018 in the EnrichR platform.

| **Following analysis by TRRUST version 2 (using both mice and human databases):** | | |
| --- | --- | --- |
| Term | Overlap | P-value |
| **RBL2_mouse** | 2/10 | **0.005623019** |
| **XBP1_human** | 2/19 | **0.019958998** |
| NRF1_mouse | 2/22 | 0.026359655 |
| PAX2_human | 1/6 | 0.067337419 |
| PBX2_human | 1/6 | 0.067337419 |
| KLF10_human | 1/7 | 0.078112883 |
| VDR_human | 2/42 | 0.084651968 |
| TBX1_mouse | 1/8 | 0.088764388 |
| DNMT3A_mouse | 1/8 | 0.088764388 |
| SP4_mouse | 1/8 | 0.088764388 |
| KCNIP3_mouse | 1/8 | 0.088764388 |
| **Using KEGG 2019 Database in EnrichR:** |  |  |
| Term | Overlap | P-value |
| **Glutamatergic synapse** | 14/114 | 5.70E-11 |
| **Long-term potentiation** | 10/67 | 4.85E-09 |
| **Endocytosis** | 17/269 | 1.66E-08 |
| **Long-term depression** | 9/61 | 3.22E-08 |
| **Calcium signaling pathway** | 14/189 | 4.42E-08 |
| **Amphetamine addiction** | 9/68 | 8.53E-08 |
| Circadian entrainment | 10/99 | 2.21E-07 |
| **Valine, leucine and isoleucine degradation** | 8/56 | 2.47E-07 |
| **Huntington disease** | 13/192 | 3.83E-07 |
| **Dopaminergic synapse** | 11/135 | 4.90E-07 |
| Insulin secretion | 9/86 | 6.65E-07 |
| **Alzheimer disease** | 12/175 | 9.47E-07 |
| Retrograde endocannabinoid signaling | 11/150 | 1.40E-06 |
| Bacterial invasion of epithelial cells | 8/74 | 2.19E-06 |
| Gastric acid secretion | 8/74 | 2.19E-06 |
| Aldosterone synthesis and secretion | 9/102 | 2.82E-06 |
| Endocrine and other factor-regulated calcium reabsorption | 7/55 | 3.17E-06 |
| **Nicotine addiction** | 6/40 | 6.15E-06 |
| Butanoate metabolism | 5/27 | 1.29E-05 |
| **Protein processing in endoplasmic reticulum** | 10/163 | 2.05E-05 |
| Thyroid hormone synthesis | 7/73 | 2.13E-05 |
| beta-Alanine metabolism | 5/32 | 3.07E-05 |
| **Cholinergic synapse** | 8/113 | 5.07E-05 |
| Oocyte meiosis | 8/116 | 6.11E-05 |
| Alanine, aspartate and glutamate metabolism | 5/37 | 6.35E-05 |
| Fc gamma R-mediated phagocytosis | 7/87 | 6.68E-05 |
| **GABAergic synapse** | 7/90 | 8.29E-05 |
| Hippo signaling pathway | 9/159 | 9.93E-05 |
| Glycolysis / Gluconeogenesis | 6/67 | 1.23E-04 |
| **Tight junction** | 9/167 | 1.44E-04 |
| **Cocaine addiction** | 5/48 | 2.25E-04 |
| **Tryptophan metabolism** | 5/48 | 2.25E-04 |
| Fatty acid degradation | 5/50 | 2.73E-04 |
| Parkinson disease | 8/144 | 2.74E-04 |
| Salivary secretion | 6/78 | 2.86E-04 |
| Salmonella infection | 6/78 | 2.86E-04 |
| Adrenergic signaling in cardiomyocytes | 8/148 | 3.30E-04 |
| Oxytocin signaling pathway | 8/154 | 4.31E-04 |
| Gap junction | 6/86 | 4.84E-04 |
| GnRH signaling pathway | 6/90 | 6.17E-04 |

**Table S3.**

Hippocampal proteome derived from young non-Tg, aged non-Tg, and aged Tg^XBP1s^ animals. Significant alterations (*P* < 0.05) are highlighted. Table shows the genes encoding these proteins for the three groups (*n* = 4 animals/group).

**Gene**

Pfkl

Gabra1

Ctnnd2

Gnb1

Cenpv

Dlg2

Tanc2

Pgls

Lmna

Stx1b

Matr3

Cab39l

Kif5b

Hnrnph2

Actn1

Atp1a1

Baiap2

Shank3

Negr1

Cct4

Myo18a

Srcin1

Crmp1

Scrn3

Nptn

Gabrg2

Bsn

Fdxr

Psmd5

Atp2b2

Hspa4

Sdr39u1

Homer1

Cotl1

Dsp

Synpo

Nfasc

Mlf2

Lphn3

Gdi1

Gpd1

Akr7a5

Vps26b

Asrgl1

Ccdc177

Atl1

Sirpa

Prkce

Dlgap4

Rbmxl1

**Table S4.**

Enrichment analysis of the most significant proteomic alterations comparing the hippocampi of aged non Tg with aged Tg^XBP1s^ (*P* < 0.05, *n* = 4 animals/group) using the Gene Ontology 2019 database. Protein-protein networks were generated by STRING v11.

| **term description** | **observed gene count** | **background gene count** | **false discovery rate** |
| --- | --- | --- | --- |
| cellular component organization | 29 | 4560 | 1.53E-05 |
| **modulation of chemical synaptic transmission** | 10 | 367 | 1.53E-05 |
| **synapse organization** | 8 | 181 | 1.53E-05 |
| **protein localization to synapse** | 5 | 49 | 3.80E-05 |
| membrane organization | 10 | 548 | 9.35E-05 |
| generation of neurons | 15 | 1538 | 0.00021 |
| localization | 25 | 4315 | 0.00026 |
| cell morphogenesis involved in differentiation | 9 | 506 | 0.0003 |
| positive regulation of synaptic transmission | 6 | 163 | 0.0003 |
| regulation of cellular component organization | 18 | 2337 | 0.0003 |
